# Bayesian inference associates rare *KDR* variants with specific phenotypes in pulmonary arterial hypertension

**DOI:** 10.1101/2019.12.11.871210

**Authors:** Emilia M. Swietlik, Daniel Greene, Na Zhu, Karyn Megy, Marcella Cogliano, Smitha Rajaram, Divya Pandya, Tobias Tilly, Katie A. Lutz, Carrie C. L. Welch, Michael W. Pauciulo, Laura Southgate, Jennifer M. Martin, Carmen M. Treacy, Christopher J. Penkett, Jonathan C. Stephens, Harm J. Bogaard, Colin Church, Gerry Coghlan, Anna W. Coleman, Robin Condliffe, Mélanie Eyries, Henning Gall, Stefano Ghio, Barbara Girerd, Simon Holden, Luke Howard, Marc Humbert, David G. Kiely, Gabor Kovacs, Jim Lordan, Rajiv D. Machado, Robert V. MacKenzie Ross, Colm McCabe, Shahin Moledina, David Montani, Horst Olschewski, Joanna Pepke-Zaba, Laura Price, Christopher J. Rhodes, Werner Seeger, Florent Soubrier, Jay Suntharalingam, Mark R. Toshner, Anton Vonk Noordegraaf, John Wharton, Jim Wild, Stephen John Wort, NIHR BioResource for Translational Research - Rare Diseases, National Cohort Study of Idiopathic and Heritable PAH, PAH Biobank Enrolling Centers’ Investigators, Allan Lawrie, Martin R. Wilkins, Richard C. Trembath, Yufeng Shen, Wendy K. Chung, Andrew J. Swift, William C. Nichols, Nicholas W. Morrell, Stefan Gräf

**Author notes:** Corresponding authors: Dr Stefan Gräf, PhD & Professor Nicholas W. Morrell, M.D., Department of Medicine, University of Cambridge, Level 5, Cambridge University Hospitals, Box 157, Cambridge Biomedical Campus, Cambridge, CB2 0QQ, United Kingdom, /. these authors jointly supervised this work (Nicholas W. Morrell, Stefan Gräf).

## Abstract

**Background:** Approximately 25% of patients with pulmonary arterial hypertension (PAH) have been found to harbor rare mutations in disease-causing genes. To identify missing heritability in PAH we integrated deep phenotyping with whole-genome sequencing data using Bayesian statistics.

**Methods:** We analyzed 13,037 participants enrolled in the NIHR BioResource - Rare Diseases (NBR) study, of which 1,148 were recruited to the PAH domain. To test for genetic associations between genes and selected phenotypes of pulmonary hypertension (PH), we used the Bayesian rare-variant association method BeviMed.

**Results:** Heterozygous, high impact, likely loss-of-function variants in the Kinase Insert Domain Receptor (*KDR*) gene were strongly associated with significantly reduced transfer coefficient for carbon monoxide (KCO, posterior probability (PP)=0.989) and older age at diagnosis (PP=0.912). We also provide evidence for familial segregation of a rare nonsense *KDR* variant with these phenotypes. On computed tomographic imaging of the lungs, a range of parenchymal abnormalities were observed in the five patients harboring these predicted deleterious variants in *KDR*. Four additional PAH cases with rare likely loss-of-function variants in *KDR* were independently identified in the US PAH Biobank cohort with similar phenotypic characteristics.

**Conclusions:** The Bayesian inference approach allowed us to independently validate *KDR*, which encodes for the Vascular Endothelial Growth Factor Receptor 2 (VEGFR2), as a novel PAH candidate gene. Furthermore, this approach specifically associated high impact likely loss-of-function variants in the genetically constrained gene with distinct phenotypes. These findings provide evidence for *KDR* being a clinically actionable PAH gene and further support the central role of the vascular endothelium in the pathobiology of PAH.

## Introduction

Pulmonary arterial hypertension (PAH) is characterized by pulmonary vascular constriction and obliteration, causing elevation of pulmonary vascular resistance and ultimately, right ventricular failure. Molecular mechanisms such as aberrant angiogenesis^1^, metabolic reprogramming and resistance to apoptosis^2^ have been proposed to explain pulmonary vessel remodeling. A breakthrough in our understanding of the pathobiology underlying PAH was the discovery of heterozygous germline mutations in the gene encoding the bone morphogenetic protein receptor type 2 (*BMPR2*)^3^, responsible for over 70% of familial PAH (FPAH) cases and 15-20% of idiopathic PAH (IPAH) cases. A smaller proportion (up to 10%) of PAH cases are caused by mutations in activin-like kinase 1 (*ACVRL1*)^4^, endoglin (*ENG*)^5^, SMAD family member 9 (*SMAD9*)^6^, caveolin-1 (*CAV1*), involved in colocalization of BMP receptors^7^, and the potassium channel *KCNK3*, responsible for membrane potential and vascular tone^8^. We recently identified rare pathogenic variants in growth differentiation factor 2 (*GDF2*), which encodes BMP9, a major ligand of the BMPR2/ALK1 receptor complex, as well as rare variants in ATPase 13A3 (*ATP13A3*), aquaporin 1 (*AQP1*) and SRY-box 17 (*SOX17*), and reported a list of additional putative genes potentially contributing to the pathobiology of PAH^9^. Together, the established genes explain approximately 25% of cases with IPAH, allowing their reclassification as heritable PAH (HPAH) cases. To identify additional genes harboring potentially causal rare variants in IPAH cases, we increased the cohort size^10^ and deployed a recently developed Bayesian methodology (BeviMed)^11^ that incorporates phenotypic data to increase the power to detect rare risk variants.

## Methods

### Data availability

The data of the NBR study have been deposited in the European Genome-Phenome Archive^10^. The data from the US PAH Biobank (PAHBB) and the Columbia University Medical Center (CUMC) are available via an application^12^.

### Study design, ethics, and subject recruitment

The NBR was established to identify genetic causes, streamline molecular diagnosis and develop new treatments for rare diseases through whole-genome sequencing (WGS) and deep phenotyping^10^. The study comprised 18 domains (<u>Table I</u> in the Data Supplement), including 1,148 adult and pediatric subjects recruited to the PAH domain. Included were individuals diagnosed with either IPAH or HPAH, pulmonary veno-occlusive disease (PVOD) or pulmonary capillary haemangiomatosis (PCH) and a small number of healthy relatives. Recruitment of PAH cases was carried out across the nine pulmonary hypertension (PH) specialist centers in the UK and by European collaborators (see <u>Data Supplement</u>). Patients recruited to the NBR study provided informed consent for genetic analysis and clinical data capture (REC REF: 13/EE/0325); patients recruited by European collaborators consented to genetic testing and clinical data collection locally.

Patients with rare diseases recruited to domains other than PAH were used as non-PAH controls in the genetic analysis (<u>Table I</u> in the Data Supplement).

For validation, we used the US PAHBB cohort comprising exome sequencing data from 2,572 subjects diagnosed with group 1 PAH^12^ and a biobank of 440 PAH patients established at CUMC^13^.

### Phenotyping of patients

#### Clinical phenotyping and case-control cohort using phenotypic ‘tags’

Pseudonymized results of routine clinical tests were stored in the electronic data capture system *OpenClinica* (<u>Table II</u> in the Data Supplement). All cases were diagnosed between 2002 and 2017 according to international guidelines using a multidisciplinary assessment that included echocardiography, comprehensive blood testing, pulmonary function testing, overnight oximetry, isotope perfusion scanning, high-resolution computed tomography, and right heart catheterization. To aid data analysis and improve data quality, a number of quality assurance procedures were introduced (see <u>Data Supplement</u>). Diagnosis was ascertained in all cases. (<u>Figure II in the Data Supplement</u>). The full set of tags, with corresponding numbers of cases, controls, and excluded relatives, is shown in <u>Table 1</u> and further discussed in the <u>Data Supplement</u>.

**Table 1.** Definitions of labels and the number of unrelated cases and controls in the rarevariant association analysis with BeviMed. See paragraph on “Number of PAH domain samples in the analysis” in the <u>Data Supplement</u> for more details. Abbreviations: mPAP -mean pulmonary artery pressure, PH - pulmonary hypertension, PAH - pulmonary arterial hypertension, I/HPAH - Idiopathic/Hereditary Pulmonary Arterial Hypertension, PVOD - Pulmonary veno-occlusive disease, PCH - Pulmonary capillary haemangiomatosis, APAH - Associated Pulmonary Arterial Hypertension, CHD - Congenital Heart Disease, CTD - Connective Heart Disease, LHD - Left Heart Disease, LD - Lung Disease, CTEPH - Chronic Thromboembolic Pulmonary Hypertension, KCO - transfer coefficient for carbon monoxide.

| Tag | Tag description | Cases | Controls | Excluded relatives |
| --- | --- | --- | --- | --- |
| **PH** | Individuals with mPAP > 25 mmHg | 1112 | 9134 | 2786 |
| **PAH** | Patients with one of the following diagnoses: IPAH, HPAH, PVOD, PCH, APAH:CHD-PAH, APAH:CTD-PAH, APAH:HIV-PAH, APAH:PH-PAH | 1085 | 9134 | 2786 |
| **I/HPAH** | Patients with a clinical diagnosis of IPAH or HPAH | 1036 | 9134 | 2786 |
| **IPAH** | Patients with a clinical diagnosis of IPAH | 972 | 9134 | 2785 |
| **HPAH** | Patients with a clinical diagnosis of HPAH | 67 | 9136 | 2779 |
| **PVOD/PCH** | Patients with a clinical diagnosis of PVOD/PCH | 20 | 9136 | 2778 |
| **I/HPAH/PVOD/PCH** | Patients with one of the following diagnoses: IPAH, HPAH, PVOD, PCH | 1056 | 9134 | 2786 |
| **FPAH** | Patients with one of the following diagnoses: IPAH, HPAH, PVOD, PCH and a positive family history | 80 | 9136 | 2781 |
| **APAH** | Patients with one of the following diagnoses: APAH:CHD-PAH, APAH:CTD-PAH, APAH:HIV-PAH, APAH:PH-PAH | 29 | 9136 | 2778 |
| **APAH: CHD-PAH** | Patients with PAH associated with congenital heart disease | 17 | 9136 | 2778 |
| **APAH: CTD-PAH** | Patients with PAH associated with connective tissue disease | 10 | 9136 | 2778 |
| **APAH: PoPH** | Patients with PAH associated with portopulmonary hypertension | 1 | 9136 | 2778 |
| **APAH: HIV-PAH** | Patients with PAH associated with HIV | 1 | 9136 | 2778 |
| **PH-LHD** | Patients with pulmonary hypertension associated with left heart disease (Group 2) | 7 | 9136 | 2778 |
| **PH-LD** | Patients with pulmonary hypertension associated with lung disease (Group 3) | 8 | 9136 | 2778 |
| **CTEPH** | Chronic thromboembolic pulmonary hypertension (Group 4) | 6 | 9136 | 2778 |
| **PH-multifactorial** | Multifactorial pulmonary hypertension (Group 5) | 6 | 9136 | 2778 |
| **young age** | Lower age tertile (<40.8 years) | 378 | 9136 | 2785 |
| **middle age** | Middle age tertile (40.8 - 58.6 years) | 376 | 9134 | 2779 |
| **old age** | Higher age tertile (>58.6 years) | 355 | 9136 | 2778 |
| **low KCO** | KCO <50% pred. | 152 | 9136 | 2778 |
| **KCO lower tertile** | KCO <60% pred. | 211 | 9136 | 2778 |
| **KCO middle tertile** | KCO 60-80% pred. | 215 | 9136 | 2778 |
| **KCO higher tertile** | KCO >80% pred. | 215 | 9134 | 2779 |

#### Analysis of computerized tomography (CT) scans

Diagnostic chest CT scans were performed in 613 study participants and the reports were transcribed to electronic case report forms. Of the 613 scans, 294 were available for more detailed analysis. The scans were reviewed by two independent cardiothoracic radiologists at the University of Sheffield with expertise in pulmonary hypertension (AS, SR), who were blinded to the underlying diagnosis, mutation and smoking status. For consistency and reproducibility, all measurements were reported on a customized proforma. Twenty-two scans were scored by both radiologists with good interobserver agreement (for further details see <u>Data Supplement, Table III</u> and <u>IV</u>).

### Genetic association between rare variants and selected diagnostic and phenotypic tags

A schematic of the analysis pipeline is depicted in <u>Figure 1A</u>. We hypothesized that phenotypically homogenous groups of patients would also share a similar genetic etiology. In particular, we focused on the current diagnostic classification of pulmonary hypertension and stratification by age at diagnosis and KCO (% predicted), as discussed in <u>Data Supplement</u>, to define a set of phenotypic tags (<u>Table 1</u>). We defined cases as individuals carrying a particular tag whereas the individuals from the non-PAH domains served as controls (<u>Figure 1, Table I</u> in the Data Supplement). Rare variants were extracted from each gene as previously described using the WGS data^10^ (see <u>Data Supplement</u>). The analysis was restricted to canonical transcripts as annotated by Ensembl. The Bayesian model comparison method BeviMed^11^ was applied to the rare variants from a set of unrelated individuals to estimate posterior probabilities of gene-tag associations under dominant and recessive modes of inheritance. Importantly, since we selected rare variants that do not correlate strongly with coarse population structure, we did not correct for ethnicity. The association testing was only performed on unrelated individuals (<u>Table 1, Table I</u> and <u>V</u> in the Data Supplement). Patients with rare deleterious variants in previously established PAH disease genes (*BMPR2, ACVRL1, ENG, CAV1, SMAD1, SMAD4, SMAD9, KCNK3, EIF2AK4, TBX4, AQP1, ATP13A3, GDF2, SOX17*) that were deemed disease-causing by a genetic multidisciplinary team according to the ACMG Standards and Guidelines^14^ were excluded from the association testing for other genes to minimize false-positive associations. To increase power in scenarios where only variants of particular consequence types were associated with the disease risk, association models were fitted to different subsets of variants according to the impact categories defined by Ensembl (high, moderate and the combination of both; see <u>Data Supplement</u> for more details). The prior probability of association across all association models was set to 0.001. Our choice of prior was based on the assumption that approximately 30 genes with rare variants of moderate and high impact might be involved in the pathogenesis of PAH out of the 32,606 protein-coding and non-coding genes (defined by the selected gene biotypes provided by Ensembl, see <u>Data Supplement</u>). Rather than relying on significance thresholds (commonly used in frequentist approaches but discouraged in Bayesian statistical frameworks) we ranked our findings according to the posterior probability truncating the list around the level of the latest previously reported PAH risk genes.

**Figure 1.**
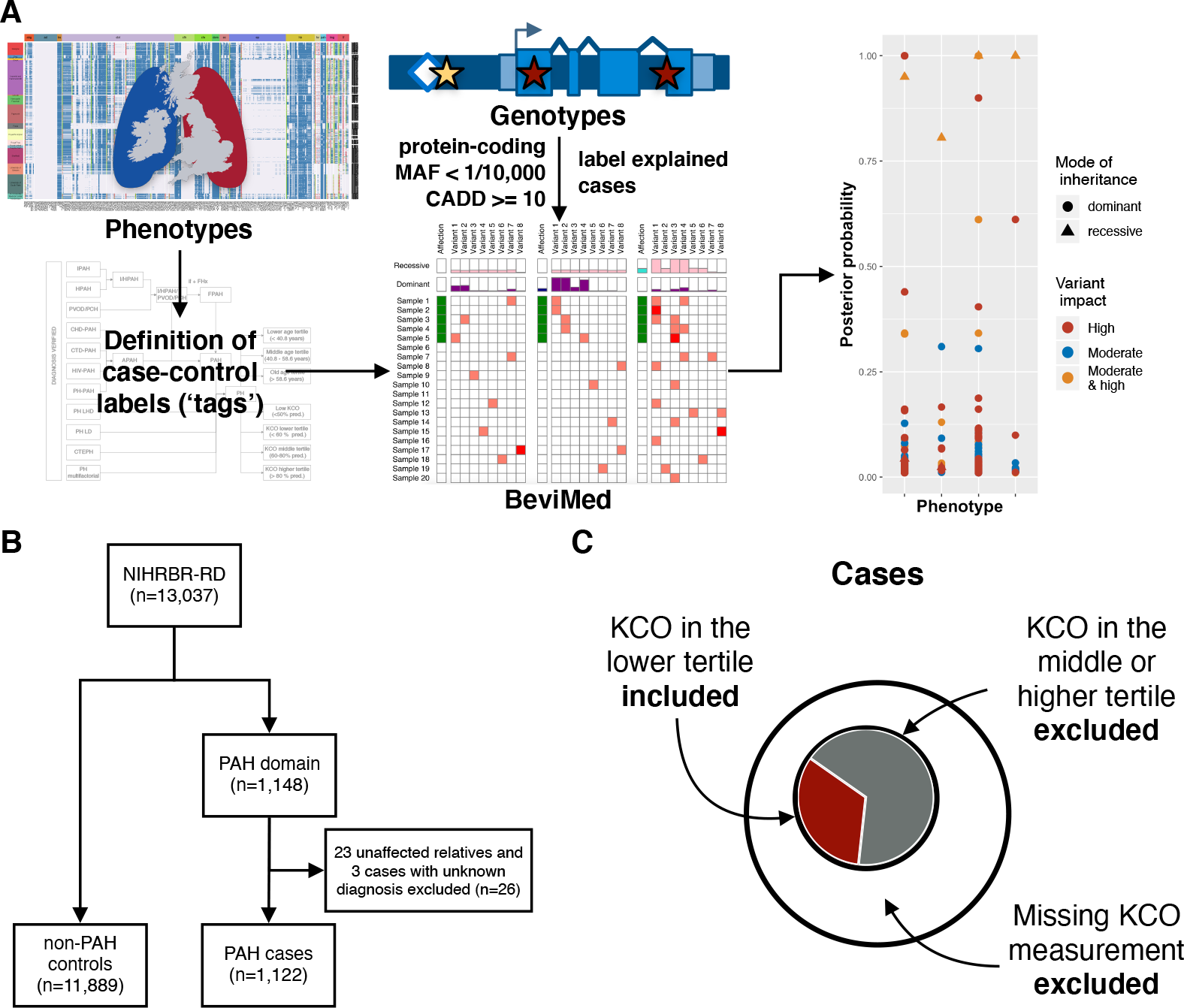
Design of the genetic association study. **A**, Overview of the analytical approach. Using deep phenotyping, data tags were assigned to patients who shared phenotypic features (for more details see <u>Figure II</u> in the Data Supplement). Rare sequence variants, called from whole-genome sequencing data, were filtered and explained cases were labeled. BeviMed was applied to a set of unrelated individuals to estimate the posterior probability of gene-tag associations. **B**, Consort diagram summarizing the size of the study cohort. **C**, Schematic representation of the definition of cases, exemplified by the KCO lower tertile tag. Cases were defined as individuals carrying a particular tag, whereas patients with missing information or those without a tag were removed from the gene-tag association testing. Individuals from non-PAH domains served as controls. KCO - transfer coefficient for carbon monoxide, MAF - minor allele frequency.

## Results

### Characterization of study cohorts and tag definition

Whole-genome sequencing was performed in 13,037 participants of the NBR study, of which 1,148 were recruited to the PAH domain^10^. The PAH domain included 23 unaffected parents and three cases with an unknown phenotype, which were removed from the analysis (<u>Figure 1B</u>). Of the remaining 1,122 participants, 972 (86.6%) had a clinical diagnosis of IPAH, 73 (6.5%) of HPAH, and 20 (1.8%) were diagnosed with PVOD/PCH. Diagnosis verification revealed that 57 participants (5%) had a diagnosis other than IPAH, HPAH or PVOD/PCH. These cases were subsequently relabelled and moved to the respective tag group for analysis. The comprehensive clinical characterisation of the study cohort is shown in <u>Table VI</u> in the Data Supplement. In summary, the median age at diagnosis was 49[35;63] years with a female predominance of 68%. Europeans constituted 84% of the study cohort. Overall survival in the studied population was 97% at one year, 91% at three years and 84% at five years. As expected, there was a significant difference in survival between prevalent and incident cases. In prevalent cases, survival at one, three, and five years was 98%, 93%, and 87%, whereas in incident cases it was 97%, 84%, and 72%, respectively. Median transfer coefficient for carbon monoxide (KCO) in the entire studied population was 71[52;86]% predicted. Cases in the lower tertile or below the KCO threshold of 50% predicted were more commonly male, older at diagnosis, had a current or past history of cigarette smoking and an increased number of cardiorespiratory comorbidities (see <u>Figure III, Table VII, VIII</u>, and <u>IX</u> in the Data Supplement). Survival in these groups was significantly worse than in those with preserved or mildly reduced KCO (<u>Figure III A-D</u> in the Data Supplement). After adjusting for confounding factors (age, sex, comorbidities, smoking status and whether the case was prevalent or incident), KCO remained an independent predictor of survival (<u>Table X</u> in the Data Supplement).

Age at diagnosis was calculated as age at the time of diagnostic right heart catheterisation and was available in all but 10 cases. Patients in the higher age tertile showed more functional impairment despite milder hemodynamics, lower FEV1/FVC ratio and KCO [% predicted], as well as mild emphysematous and fibrotic changes on CT scans (Figure III E and F and <u>Table XI</u> in the Data Supplement).

### Rare variants in previously established genes

We identified variants in previously established genes (namely, *BMPR2, ACVRL1, ENG, SMAD1, SMAD4, SMAD9, KCNK3, TBX4, EIF2AK4, AQP1, ATP13A3, GDF2, SOX17*) in 271 (24.2%) of the 1,122 cases and interpreted them based on the ACMG standards and guidelines^14^. The majority of these variants have already been described in Gräf *et al*.^*9*^ (see <u>Data Supplement</u>).

### Rare variant association testing

We used BeviMed to consolidate previously reported PAH genes and to discover novel genotype-phenotype associations. Of note, cases explained by rare deleterious variants in previously established genes were only included for the association testing with the respective disease gene (see Methods). This analysis identified 40 significant gene-tag associations with posterior probability (PP) above 0.75 (<u>Table 2</u> and <u>Figure 2A</u>). *BMPR2, TBX4, EIF2AK4, ACVRL1* and *AQP1* showed the highest association (PP≥0.99) but we also confirmed significant associations in the majority of other previously identified genes. Individuals with rare variants in *BMPR2, TBX4* (high impact), *EIF2AK4* (biallelic) and *SOX17* had a significantly younger age of disease onset (tag: young age). We also confirmed the association of rare variants in *AQP1* with FPAH (log(BF)=10.023, PP=0.958). The refined phenotype approach corroborated the association between high impact variants in *BMPR2* and preserved KCO (KCO higher tertile, log(BF)=99.923, PP=1) together with an association of biallelic *EIF2AK4* mutations with significantly reduced KCO (KCO <50% predicted, log(BF)= 29.741, PP=1).

**Table 2.**
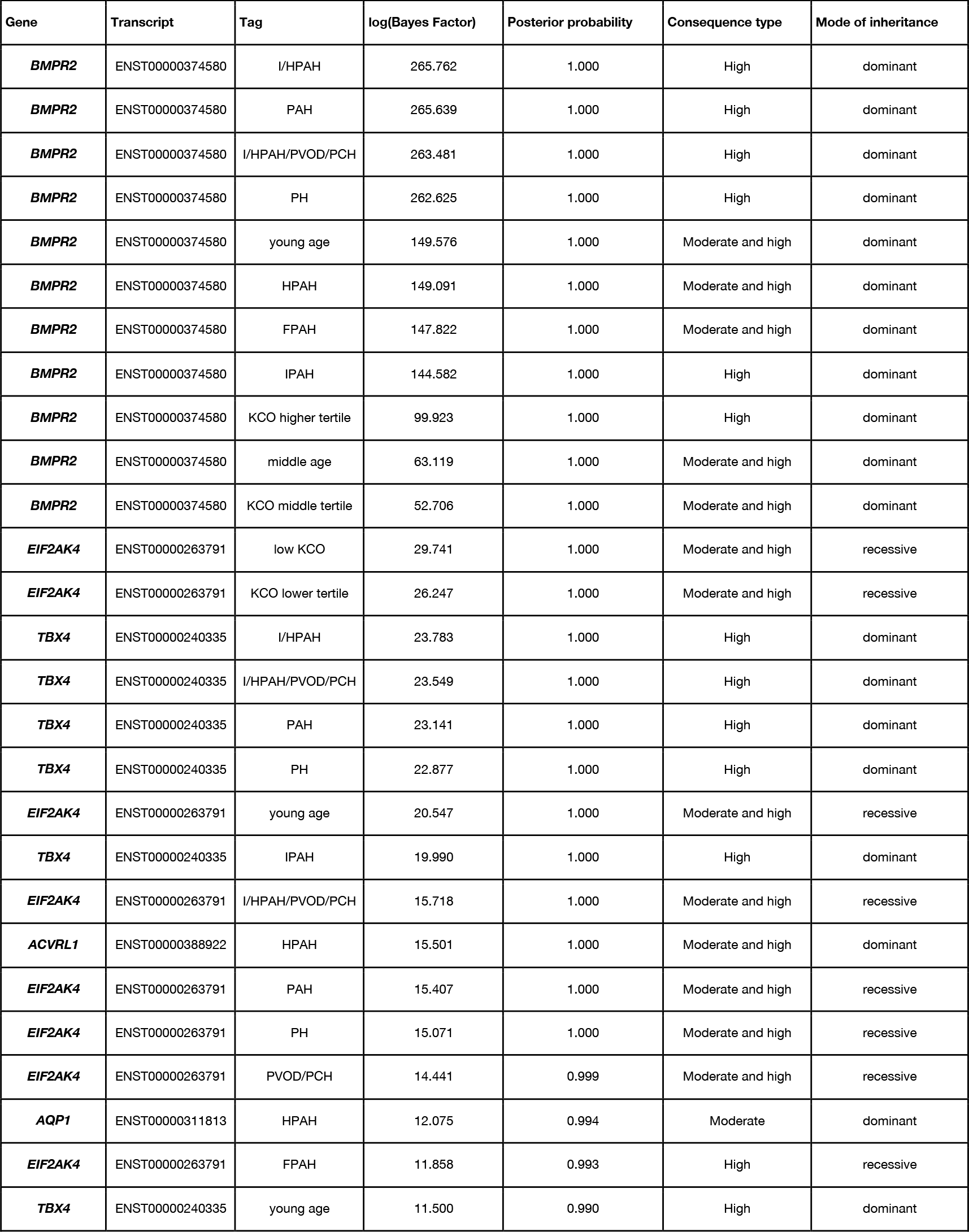
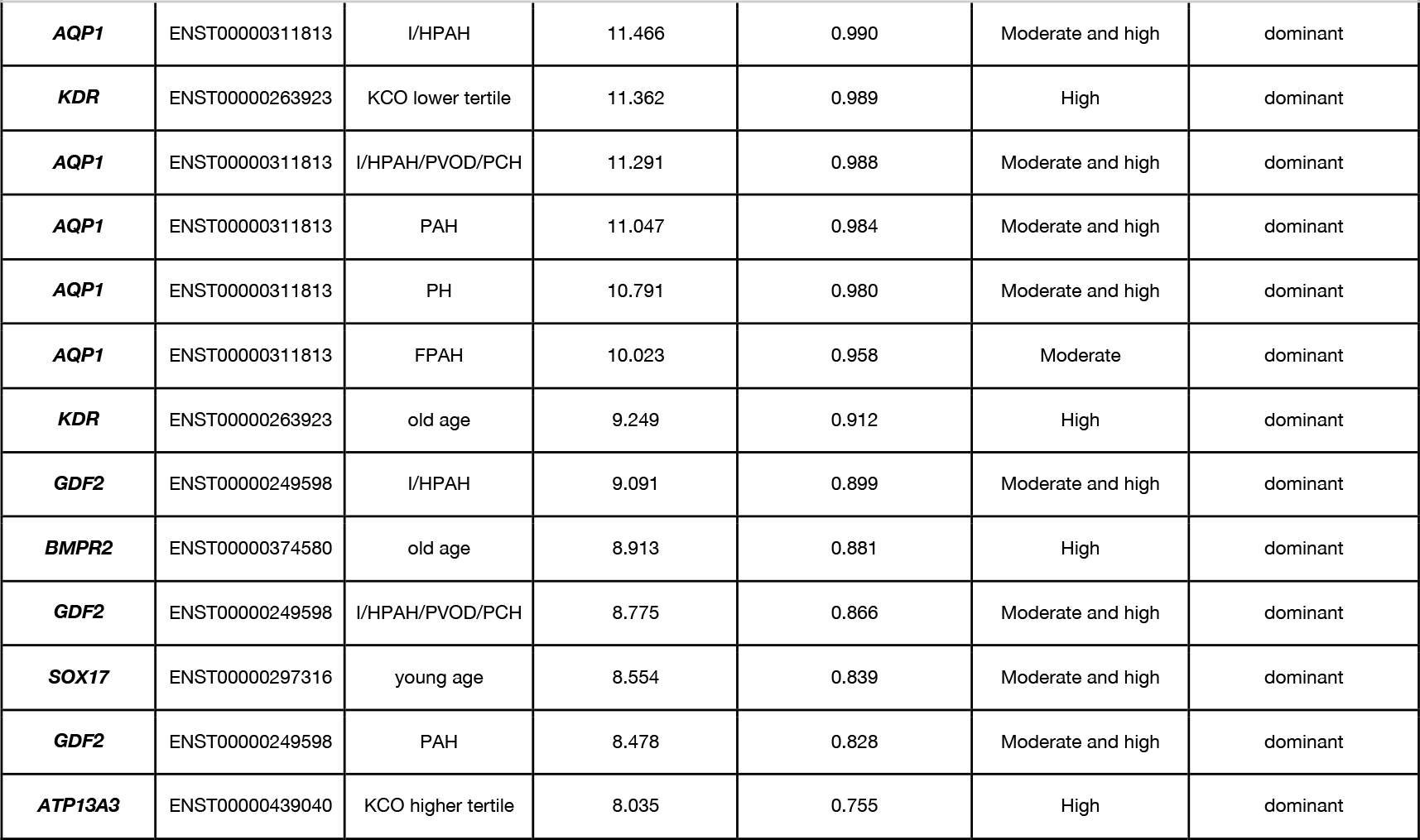
BeviMed analysis results. Posterior probabilities and Bayes Factors (BF) of gene-tag associations. The BF is the ratio between the probabilities of the data under H1 and under H0. The observed data are BF times more likely under H1 than under H0, and so the larger the BF, the stronger the support in the data for H1 compared with H0. The “High” category comprises only variants of high impact, including loss-of-function variants and large deletions; the “Moderate” category contains variants of moderate impact, including missense variants or variants of consequence type “non_coding_transcript_exon_variant”; the combined category “Moderate and High” includes both respective consequence types.

**Figure 2.**
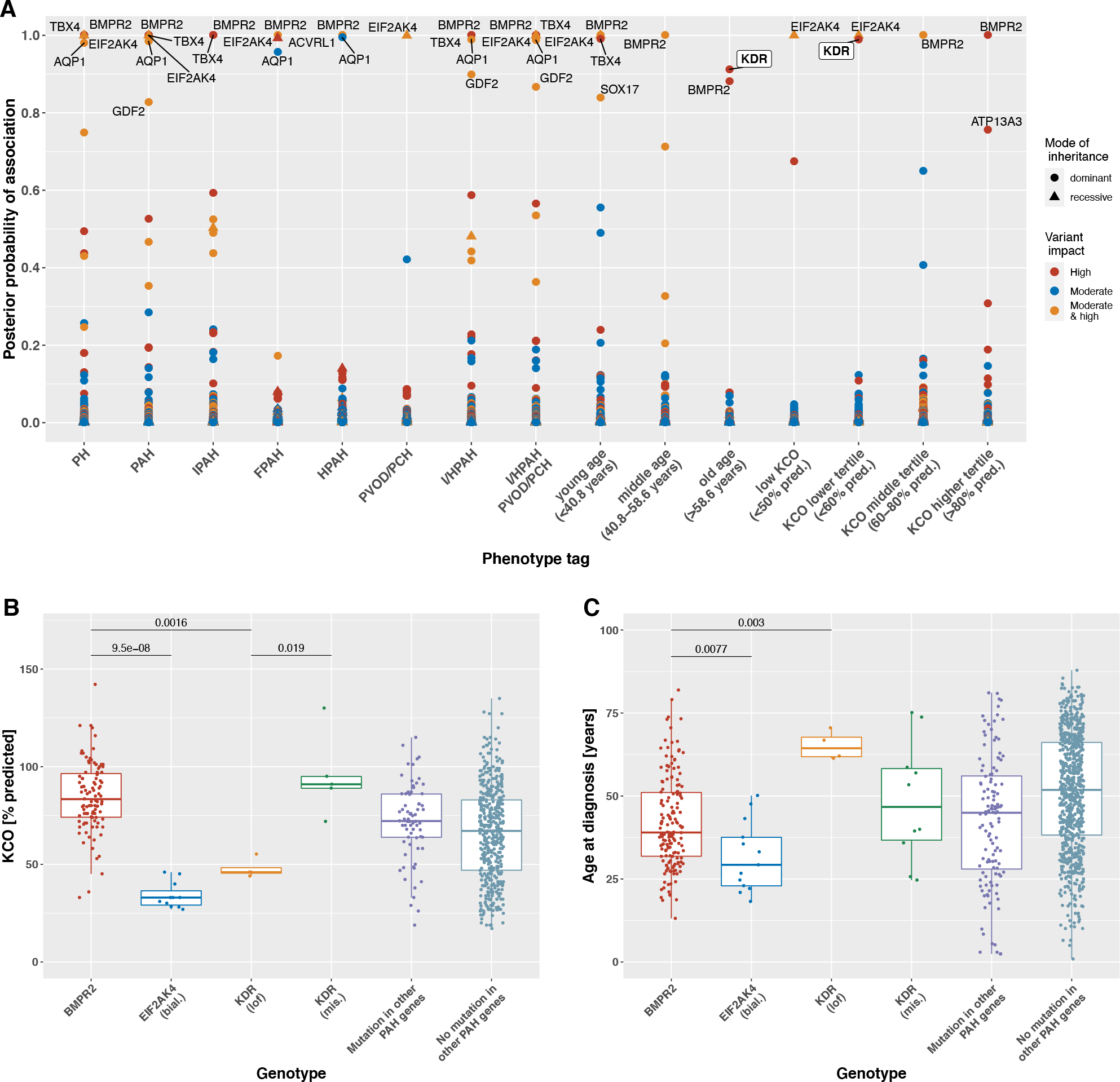
Rare variant association study results revealing established and novel genotype-phenotype links. **A**, Figure showing phenotype tags on the x-axis and corresponding posterior probability of genotype-phenotype association on the y-axis, as calculated by BeviMed. The definitions of the tags are listed in <u>Table 1</u>. Shape and colour of points indicate the mode of inheritance and impact/consequence type of variants driving the association. Box-and-whisker plots showing the distribution of (**B**) the transfer coefficient for carbon monoxide (KCO) and (**C**) the age at diagnosis stratified by genotype across the PAH domain. The two-tailed Wilcoxon signed-rank test was used to determine differences in the medians of the distributions, which are indicated by the bars at the top of the figures providing the respective p-values. Abbreviations: bial. - biallelic, lof - loss-of-function, mis. - missense.

Under an autosomal dominant mode of inheritance, high impact variants in the Kinase Insert Domain Receptor (*KDR)* were associated with a significantly reduced KCO (KCO lower tertile, log(BF)=11.362, PP=0.989) and older age at diagnosis (tag: old age, log(BF)=9.249, PP=0.912).

### Rare high impact variants in the new PAH candidate gene *KDR*

We identified five ultra-rare high impact variants in *KDR* in the study cohort. Ultra-rare variants exist in the general population only at a frequency of less than 1 in 10,000 (0.01%). Four of these were in PAH cases: one frameshift variant in exon 3 of 30 (c.183del, p.Tryp61CysfsTer16), two nonsense variants, one in exon 3 (c.183G>A, p.Trp61Ter) and one in exon 22 (c.3064C>T, p.Arg1022Ter) and one splice acceptor variant in intron 4 of 29 (c.490- 1G>A). In addition, one nonsense variant was identified in exon 27 (p.Glu1206Ter) in a non-PAH control subject (<u>Table 3</u>). This latter nonsense variant appears late in the amino acid sequence, in exon 27 of 30, and hence is likely to escape nonsense-mediated decay, but this remains to be studied functionally. Furthermore, 13 PAH cases (1%) and 108 non-PAH controls (0.9%) harbored rare, predicted-deleterious *KDR* missense variants of moderate impact (<u>Figure 3</u>). The missense variant carriers, however, did not exhibit a reduced KCO or older age at diagnosis. Instead, these patients show the opposite trend in KCO (<u>Figure 2B and C</u>). Importantly, seven of the 13 *KDR* missense variants seen in PAH cases were also detected in several non-PAH controls and thus, are of unknown significance. Furthermore, three of these missense variants co-occurred with a predicted-deleterious variant in an established PAH risk gene (two patients carried also a variant in *BMPR2* and one a variant in *AQP1*).

**Table 3.**
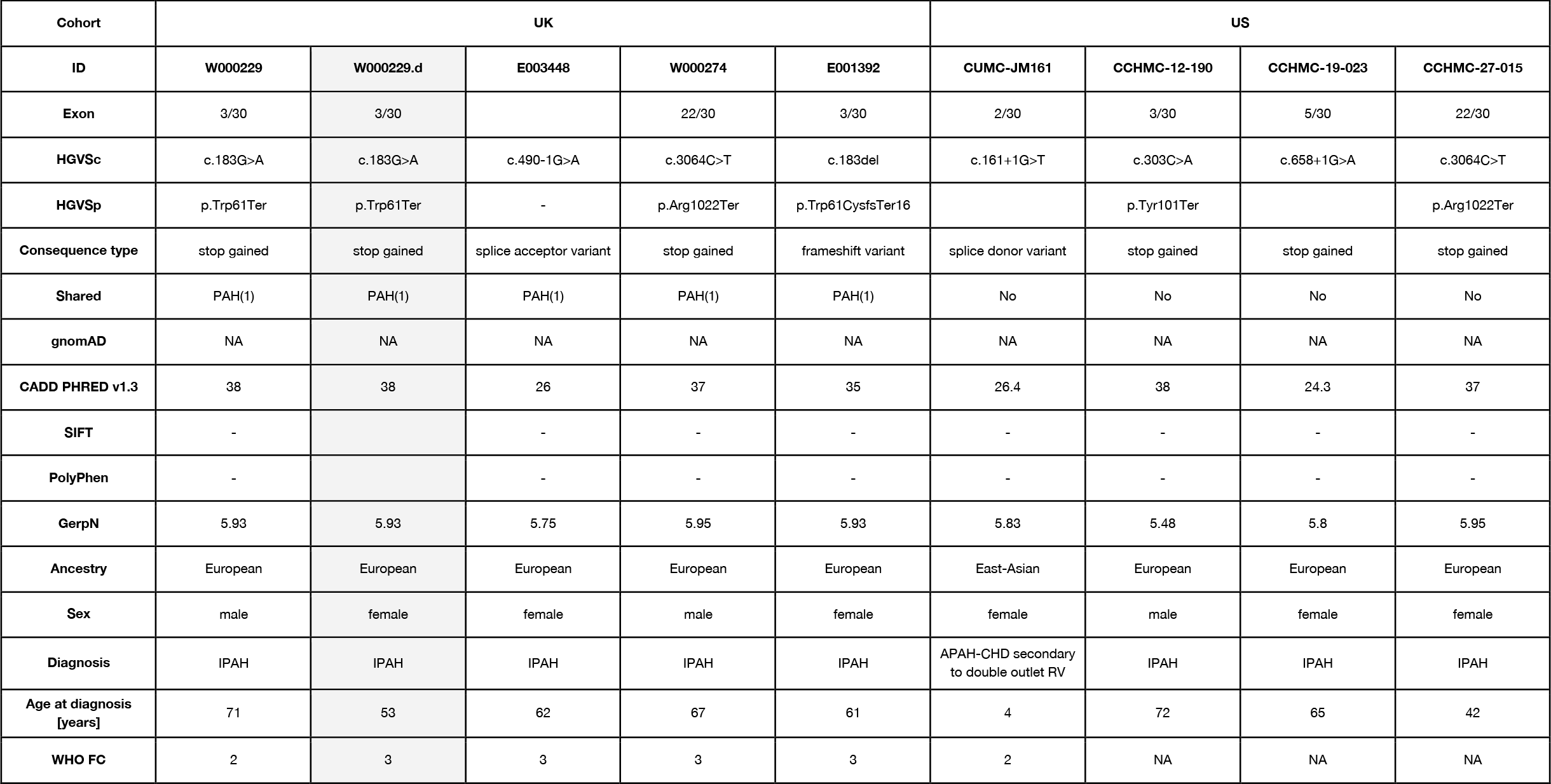
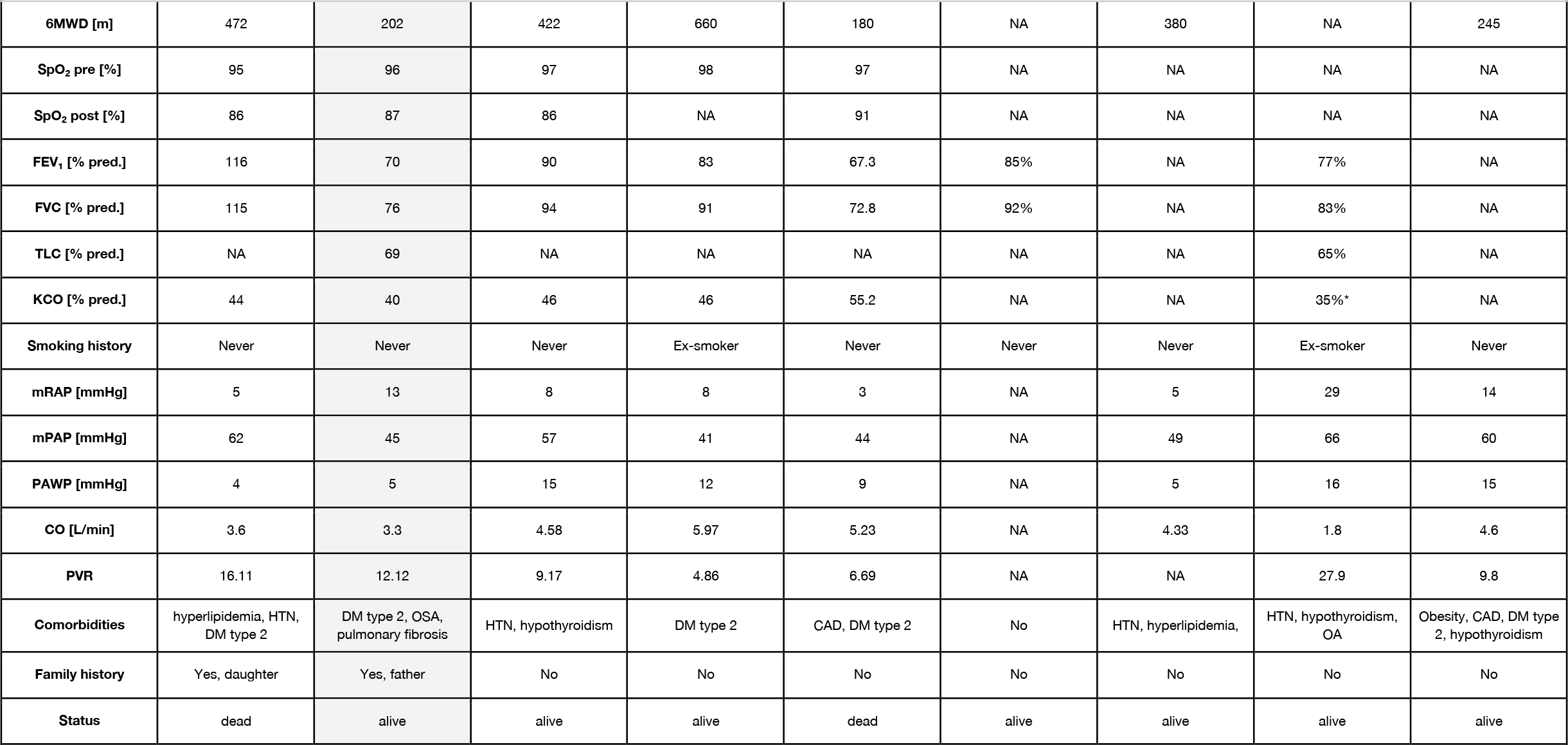
Gene changes for IPAH patients harboring likely loss-of-function variants in the *KDR*gene. None of the *KDR* variants have previously been reported in gnomAD, ExAC or internal controls. HGVSc notations are based on transcript sequence ENST00000263923.4. HGVSp notations are based on the amino acid sequence ENSP00000263923.4. Abbreviations: *KDR*- Kinase insert domain receptor, WHO FC - World Health Organization functional class,6MWD - 6-minute walk distance, SpO_2_ - arterial oxygen saturation, mRAP - mean right atrial pressure, mPAP - mean pulmonary artery pressure, mPAWP - mean pulmonary artery wedge pressure, CO - cardiac output, PVR - pulmonary vascular resistance, FEV_1_ - forced expiratory volume in 1 sec, FVC - forced vital capacity, KCO - transfer factor coefficient for carbon monoxide, HTN - systemic hypertension, CAD - coronary artery disease, OA - osteoarthritis, DM - diabetes mellitus, OSA - obstructive sleep apnea, ASD - atrial septal defect, VSD - ventricular septal defect. * DLCO % predicted.

**Figure 3.**
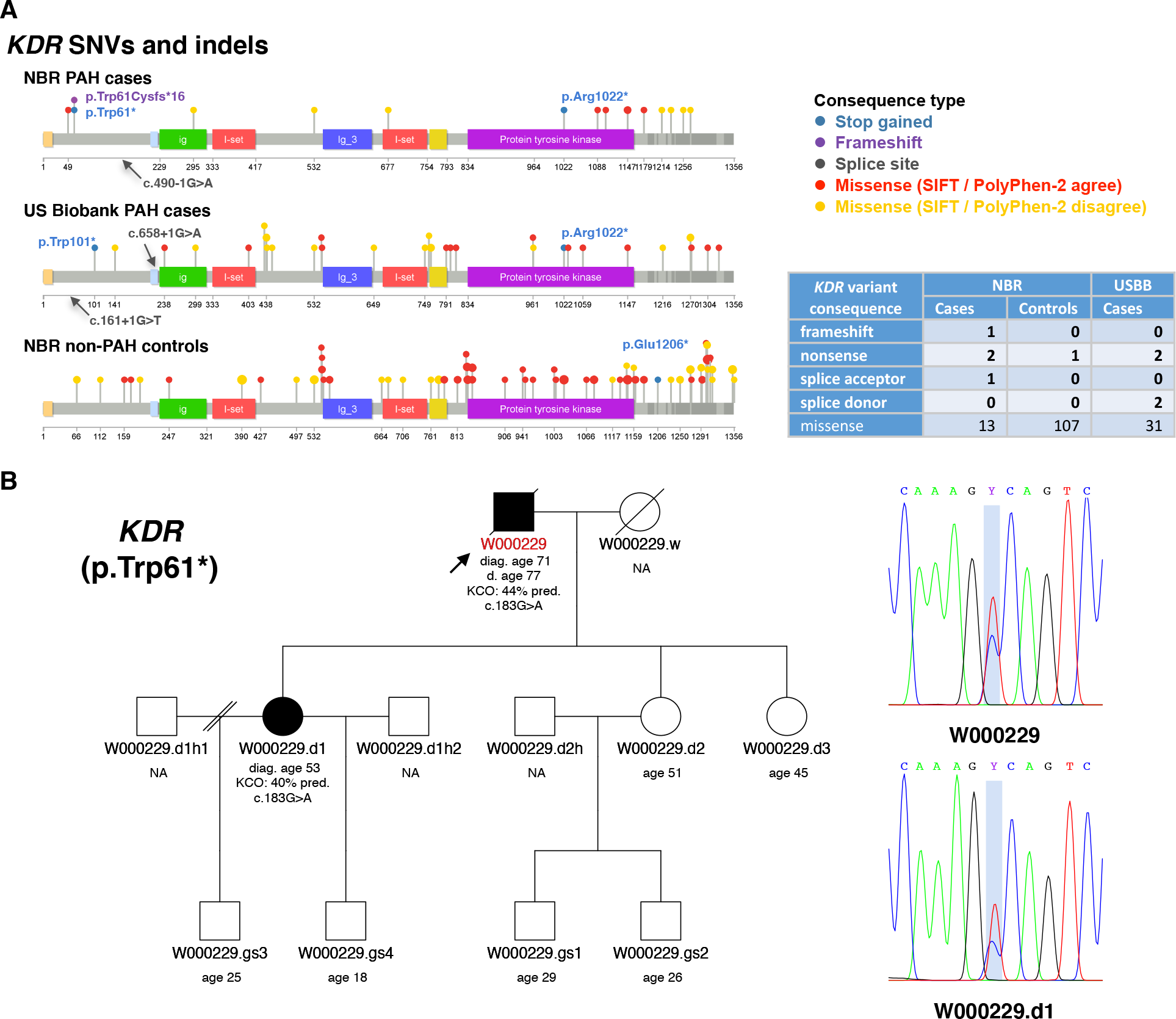
Summary of rare single nucleotide variants (SNVs) and small insertions and deletions (indels) identified in the novel PAH candidate gene *KDR*. **A**, Only rare predicted deleterious variants in *KDR* are shown (MAF<1/10,000 and CADD≥10). SNVs and indels are represented by colored lollipops on top of the protein sequence. The domain annotations were retrieved from Uniprot (accession number P35968). Lollipop colors indicate the consequence type and sizes represent the variant frequency within a cohort. Missense variants that are predicted to be deleterious (SIFT) and damaging (PolyPhen-2) are colored in red, otherwise in yellow (i.e. SIFT and PolyPhen-2 disagree). High impact variants are labelled with the respective HGVS notation. The number of variants by predicted consequence type and cohort is provided in the table. **B**, Familial segregation of *KDR* nonsense variant c.183G>A (p.Trp61*) with PAH (i.e. reduced KCO and late onset) from father (W000229) to daughter (W000229.d). Sanger sequencing results are shown in the chromatograms.

### Clinical characterization of *KDR* mutation carriers

Patients with high impact variants in *KDR* were older and exhibited significantly reduced KCO similar to biallelic *EIF2AK4* mutation carriers and in contrast to *KDR* missense variant and *BMPR2* mutation carriers (<u>Figure 2B and C</u>). Three of the four cases did not have a history of smoking. CT scans for all four patients showed a range of mild lung parenchymal changes (<u>Figure 4</u>). W000229 had evidence of mild mainly subpleural interstitial lung disease (ILD), mild emphysema, and air trapping. W000274 had signs of ILD with traction bronchiectasis in the lower zones, mild air trapping, and mild diffuse ground-glass opacities (GGO) and neovascularity. E001392 showed mild centrilobular GGO in addition to moderate pleural effusion and a trace of air trapping, but no ILD. In these cases, it seemed likely that the observed parenchymal changes contributed to the low KCO. In contrast, E003448 had a low KCO despite only a trace of central nonspecific GGO on the CT images. Comparisons of CT findings between patients harboring deleterious mutations in *BMPR2, EIF2AK4, KDR*, other PAH risk genes and patients without mutations are presented in <u>Table IV</u> in the Data Supplement. There were no differences in the frequency of comorbidities between patients harboring missense and loss-of-function variants in *KDR* although the frequency of systemic hypertension was high (44%) (<u>Table XII</u> in the Data Supplement). Survival analysis could not be conducted due to the small number of mutation carriers, as well as only two events occurring in this group. Following the death of W000229, his daughter, aged 53, was diagnosed with PAH and had a reduced KCO at 40% predicted. On the CT scan, mild interstitial fibrosis was observed (<u>Figure V</u> in the Data Supplement). Sanger sequencing confirmed that father and daughter carried the same deleterious *KDR* nonsense variant p.Trp61Ter (<u>Figure 3B</u>).

**Figure 4.**
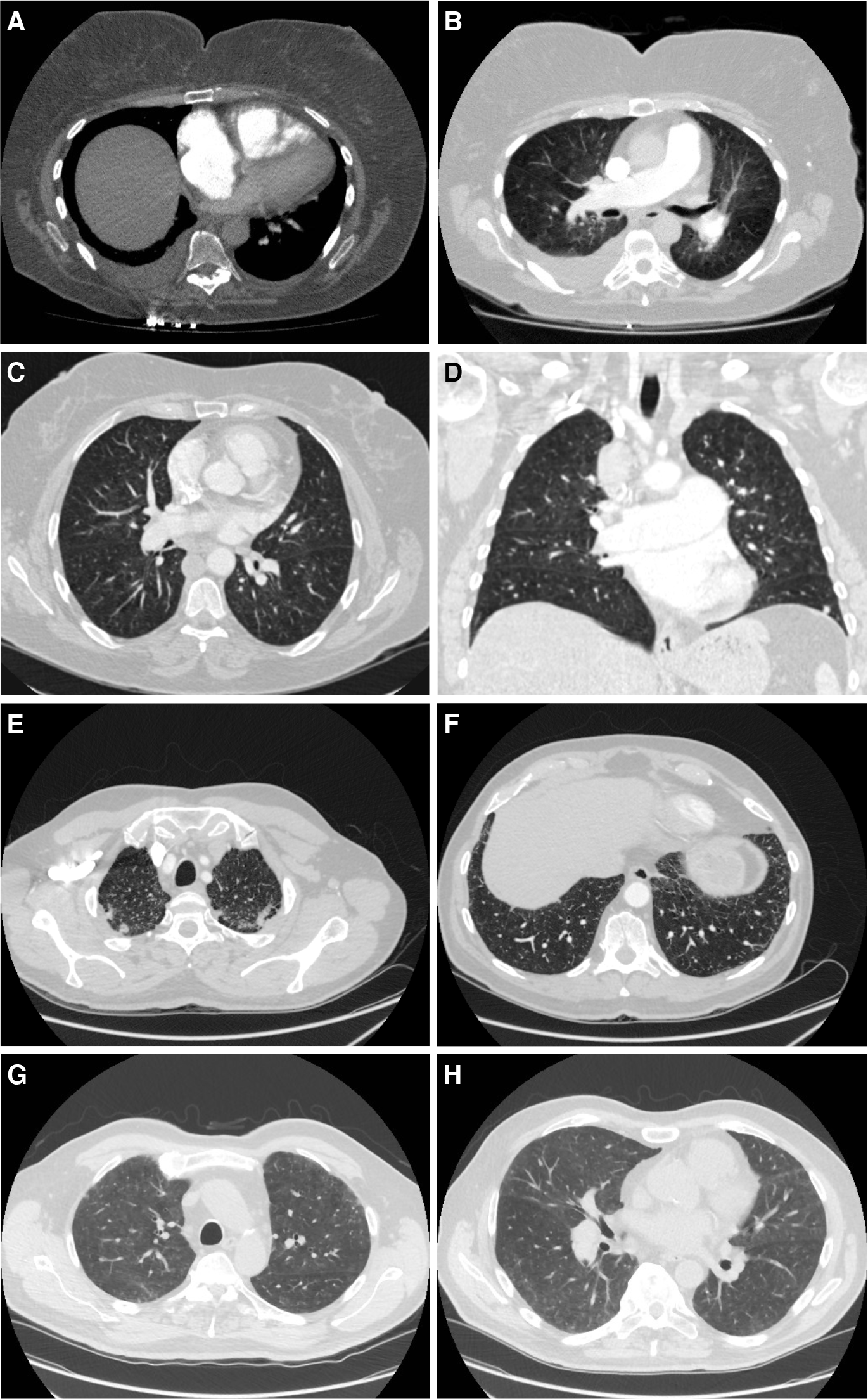
Chest computerized tomography (CT) scans of patients carrying high impact *KDR* mutations. **A**, Axial image of pulmonary CT angiogram at the level of the right ventricle (RV) moderator band, showing flattening of interventricular septum, leftwards bowing of the interatrial septum and the enlargement of the right atrium (RA) and RV, indicative of RV strain; bilateral pleural effusion, larger on the right side. **B**, Axial image of a pulmonary CT angiogram demonstrating enlarged pulmonary artery and mild central lung ground-glass opacity (GGO). **C**, Axial high-resolution CT slice of the chest in the lung window showing a trace of non- specific GGO with a central distribution. **D**, Coronal image showing the trace of central GGO and enlarged central pulmonary arteries. Axial high-resolution CT slice of the chest in the lung window showing apical subpleural fibrosis (**E**), and very minor subpleural fibrosis at the lung bases (**F**). Axial high-resolution CT slice of the chest in the lung window showing subpleuralGGO at apical level (**G**), and mild GGO at mid-thoracic level (**H**). Patients: E001392 (**A, B**), E003448 (**C, D**), W000229 (**E, F**), W000274 (**G**,**H**).

### Additional *KDR* cases in US PAH cohorts

To seek further evidence for *KDR* as a new candidate gene for PAH, we analyzed subjects recruited to the PAHBB^12^ and the CUMC^13^ to identify additional patients carrying predicted pathogenic rare variants. Four additional individuals harboring rare high impact *KDR* variants were identified. These comprised, two nonsense variants, one in exon 3 (c.303C>A, p.Tyr101Ter) and one in exon 22 (c.3064C>T, p.Arg1022Ter) and two splice donor variants, one in intron 2 of 29 (c.161+1G>T) and one in intron 5 (c.658+1G>A). Interestingly, the nonsense variant p.Arg1022Ter appeared in both cohorts (<u>Figure 3</u>). Patient-level data for these individuals are summarized in <u>Table 3</u>. Three of the four patients were diagnosed with idiopathic PAH at 72, 65 and 42 years respectively, whereas one patient was diagnosed at age four with PAH associated with double outlet right ventricle. The diffusing capacity of carbon monoxide was available for one patient and was decreased at 35% predicted, with minor pleural scarring in the left upper lobe found on CT imaging. Two out of four patients (50%) harboring a high impact variant in *KDR* had been diagnosed with systemic hypertension.

## Discussion

One of the critical steps in identifying novel, causative genes in rare disorders is the discovery of genotype-phenotype associations to inform patient care and outcomes. A pragmatic focus on deeply phenotyped individuals and “smart” experimental design provides additional leverage to identify novel risk variants^15^. To deploy this approach in PAH we brought together phenotypic and genetic data using BeviMed^11^. This Bayesian framework allows the inclusion of prior information regarding the hypothesis being tested in a flexible manner and compares a range of possible genetic models in a single analysis. To generate case-control labels, we tagged PAH cases with diagnostic labels and stratified them by age at diagnosis and KCO. Analyses were then performed to identify associations between tags and ultra-rare gene variants under dominant and recessive modes of inheritance and different variant impact categories.

Our BeviMed analysis provided strong statistical evidence of an association between ultra-rare, high impact variants in *KDR* and PAH with significantly reduced KCO and older age at diagnosis under a dominant mode of inheritance. Strikingly, likely loss-of-function variants in *KDR* exist in the general population with a frequency of only 4-7 per 100,000 (see <u>Table 4</u> in the Data Supplement). In contrast, we identified four PAH cases in the NBR cohort which equates to almost 2 in 1,000. Additionally, the statistical constraint metrics provided by gnomAD^16^ strongly suggest that loss-of-function variants in *KDR* are not tolerated (pLI = 1; o/e = 0.15 (0.09 - 0.25)). Besides the statistical evidence, we also identified one additional case with a family history, which together with a recently published case report of two families in which loss-of-function variants in *KDR* segregated with PAH and significantly reduced KCO^17^, amounts to three reported familial cases with a distinct phenotype. VEGFR2, which is encoded by *KDR*^*18*^, binds VEGFA, a critical growth factor for physiological and pathological angiogenesis in vascular endothelial cells. In mice, even though VegfA haploinsufficiency is embryonically lethal^19^, heterozygosity of its receptor, Vegfr2, is compatible with life and unperturbed vascular development^20^. The role of VEGF signaling in the pathogenesis of PAH has been an area of intense interest since increased expression of VEGF, VEGFR1 and VEGFR2 were reported in rat lung tissue in response to acute and chronic hypoxia^21^. An increase in lung VEGF has also been reported in rats with PH following monocrotaline exposure^22^. In humans, VEGF-A is highly expressed in plexiform lesions in patients with IPAH^23^. In addition, inhibition of VEGF signaling by SU5416 (sugen) combined with chronic hypoxia triggers severe angioproliferative PH^24^. SU5416, a small-molecule inhibitor of the tyrosine kinase segment of VEGF receptors, inhibits VEGFR1^25^ and VEGFR2^26^ causing endothelial cell apoptosis, loss of lung capillaries and emphysema^27^. Further evidence supporting the role of VEGF inhibition in the pathobiology of PAH comes from reports of PH in patients treated with bevacizumab^28^ and the multi-tyrosine kinase inhibitors^29,30^. Mutations in *KDR* have also been linked to congenital heart diseases. Bleyl *et al*. reported that *KDR* might be a candidate for familial total anomalous pulmonary venous return^31^. In addition, haploinsufficiency at the *KDR* locus has also been associated with tetralogy of Fallot^32^. We identified one patient in the CUMC cohort with PAH associated with congenital heart disease harboring a *KDR* likely protein-truncating splice donor variant (c.161+1G>T).

**Table 4.** Comparison of high impact likely loss-of-function variants in *KDR* in the Humanlarge-scale sequencing reference populations gnomAD and TOPMed with the NBR non-PAH controls and PAH cases. Abbreviations: gnomAD - The Genome Aggregation Database, TOPMed - The Trans-Omics for Precision Medicine program, KDR - kinase insert domain receptor, NBR - NIHR BioResource - Rare Diseases, LoF - loss of function.)

| Large-scale sequencing population | High impact LoF variants in *KDR* | Individuals | Alleles | Frequency |
| --- | --- | --- | --- | --- |
| gnomAD (v2.1)* | 20 | 141,456 | 282,912 | 0.000071 |
| gnomAD (v3) | 10 | 71,702 | 143,404 | 0.000070 |
| TOPMed | 5 | 62,784 | 125,568 | 0.000040 |
| NBR non-PAH controls | 1 | 11,889 | 23,778 | 0.000042 |
| NBR PAH cases | 4 | 1,122 | 2,244 | 0.001783 |

\**KDR* constraint metrics: pLI = 1; o/e = 0.15 (0.09 - 0.25); exp(LoF) = 73; obs(LoF) = 11

In the present study, we highlight that deep clinical phenotyping, in combination with genotype data, can improve the identification of novel disease risk genes and disease subtypes. *KDR* was already identified as a possible candidate gene, which did not achieve genome-wide significance, in our previous rare variant association study^9^. In combination with deep phenotyping data, *KDR* reached in the present study a significance level comparable to the most commonly affected genes in PAH. Reduced KCO, which reflects impairment of alveolar-capillary membrane function, has been noted in the analysis of early PAH registry data^33^ to be an independent predictor of survival. Decreased KCO was also found in patients with PVOD/PCH with or without biallelic *EIF2AK4* mutations^34^. Although some reduction in KCO is one of the typical features of pulmonary vascular disease, PVOD patients show the lowest KCO values when compared to IPAH or CTEPH. In contrast, KCO is relatively preserved in *BMPR2* mutation carriers^35^. Strong association with survival and a link with other causative mutations makes the KCO phenotype particularly attractive for stratification in genetic studies.

As lung disease should always be taken under consideration as a cause of low KCO, we applied the World Symposium on PH criteria^36^ to exclude lung disease as a cause of PH: TLC ≥70% pred., FVC ≥70% pred., FEV1 ≥60% pred., and no severe fibrosis and/or emphysema on chest CT. None of the cases carrying a high impact variant in *KDR* met these criteria, although two of the four patients did show evidence of early ILD. Another potential reason for low KCO in the PAH population is the diagnosis of PVOD/PCH^37^. Careful analysis of CT scans and clinical data did not reveal convincing evidence for this diagnosis in *KDR* mutation carriers. Cigarette smoking is a well-known factor leading to the decrease of KCO. Only one of the four *KDR* high impact variant carriers had a significant 15 pack-years smoking history, but with no signs of emphysema on CT. These findings suggest that loss-of-function variants in *KDR* are associated with a form of PAH characterized by a range of lung parenchymal abnormalities, including small airways disease, emphysema and ILD, as two of the four patients harboring a high impact variant in *KDR* had mild fibrotic lung changes. Notably, patients with mutations in other PAH risk genes, or those without the identified genetic mutation, showed less than 10% incidence of fibrotic changes on CT imaging. Further larger studies are needed to determine the full range of lung parenchymal abnormalities in PAH cases with deleterious variants in *KDR*.

In this study, we have assumed that PAH is a monogenic condition, which is caused by either deleterious heterozygous or biallelic variants in a single gene. This assumption, although widely supported by the literature, may not be entirely accurate. Alternatively, some cases of PAH might represent an oligogenic inheritance involving two or more genes. Although not statistically explored in the current analysis we found a total of 22 PAH cases carrying deleterious variants in more than one PAH gene. These variants could contribute as genetic modifiers, impacting penetrance and/or expressivity. In this analysis, we have explored only a limited number of clinical phenotypes. Further studies with larger numbers of phenotypic tags derived from clinical and molecular data will increase the power to detect new associations. Finally, KCO measurements were missing for a proportion of patients which could introduce a selection bias, although all the deleterious variants in *KDR* had phenotypic data available in the UK cohort.

In summary, this study shows that deep phenotyping enables patient stratification into subgroups with shared pathobiology and with increased power to detect new genotype-phenotype associations. We provide statistical evidence for an association between high impact, likely loss-of-function variants in *KDR* and significantly decreased KCO and later disease onset, further supported by familial segregation.

## Acknowledgments

We thank NIHR BioResource volunteers for their participation, and gratefully acknowledge NIHR BioResource centers, NHS Trusts and staff for their contribution. We thank the National Institute for Health Research and NHS Blood and Transplant. The views expressed are those of the author(s) and not necessarily those of the NHS, the NIHR or the Department of Health and Social Care.

We thank the research nurses and coordinators at the specialist pulmonary hypertension centers involved in this study. We acknowledge the support of the Imperial NIHR Clinical Research Facility, the Netherlands CardioVascular Research Initiative, the Dutch Heart Foundation, Dutch Federation of University Medical Centres, the Netherlands Organization for Health Research and Development and the Royal Netherlands Academy of Sciences. We thank all the patients and their families who contributed to this research and the Pulmonary Hypertension Association (UK) for their support. We also thank Kathryn Auckland for proofreading the manuscript. We thank contributors, including the Pulmonary Hypertension Centers who collected samples used in this study, as well as patients and their families, whose help and participation made this work possible. Exome sequencing and genotyping data were generated by the Regeneron Genetics Center.

PAH Biobank Enrolling Centers’ Investigators: Russel Hirsch, MD; R. James White, MD, PhD; Marc Simon, MD; David Badesch, MD; Erika Rosenzweig, MD; Charles Burger, MD; Murali Chakinala, MD; Thenappan Thenappan, MD; Greg Elliott, MD; Robert Simms, MD; Harrison Farber, MD; Robert Frantz, MD; Jean Elwing, MD; Nicholas Hill, MD; Dunbar Ivy, MD; James Klinger, MD; Steven Nathan, MD; Ronald Oudiz, MD; Ivan Robbins, MD; Robert Schilz, DO, PhD; Terry Fortin, MD; Jeffrey Wilt, MD; Delphine Yung, MD; Eric Austin, MD; Ferhaan Ahmad, MD, PhD; Nitin Bhatt, MD; Tim Lahm, MD; Adaani Frost, MD; Zeenat Safdar, MD; Zia Rehman, MD; Robert Walter, MD; Fernando Torres, MD; Sahil Bakshi, DO; Stephen Archer, MD; Rahul Argula, MD; Christopher Barne

