## Supplemental Material for "Bayesian inference associates rare *KDR* variants with specific phenotypes in pulmonary arterial hypertension"

Swietlik EM *et al.*

### SUPPLEMENTAL MATERIAL

#### Methods

##### Subject recruitment

Recruitment of PAH cases was carried out across the nine pulmonary hypertension (PH) specialist centers in the UK and by European collaborators at the Université Paris-Saclay and Sorbonne Université (France), University of Giessen and Marburg (Germany), and hospitals in Graz (Austria), Pavia (Italy) and Amsterdam (The Netherlands).

##### Clinical data quality assurance procedures

To aid data analysis and improve data quality we introduced a number of quality assurance procedures listed below.
Data checks were written in the R language and consisted of:

1. Logical verification checks i.e. systolic/mean/diastolic pulmonary artery pressure sPAP > mPAP > dPAP; FVC [L] > FEV1 [L], CO > CI unless BSA <1
2. Checks for discrepancies between:
    1. Comorbidities and ICD10 codes using the System of Ontology-based Re-coding and Technical annotation (SORTA)[1].
    2. Comorbidities and medication i.e. comorbidity reported but lack of medication or vice versa.
    3. Reported and calculated acute vasoresponse test results.
    4. Reported lung emphysema, thromboembolic disease and fibrosis and free text search in CT reports for these abnormalities.
    5. Reported lung function abnormalities and provided numerical values.

3. Missingness assessment and handling:
    1. Twenty-four core variables (of diagnostic and prognostic significance) have been identified and a missing rate below 10% was established. Missingness tables were reported back to the centres for clarification.
    2. Subsequently, missingness in the remaining variables was assessed and checked for logical inconsistencies, i.e. PVR reported but PAWP and CO missing, CI missing while CO and BSA available.
    3. The proportion of missing data, missing data mechanisms and missing data patterns were assessed with the *vim* and *Amelia* R packages.
4. Diagnosis verification
    1. Based on the clinical information provided, a diagnosis verification algorithm was implemented in R to red-flag patients with a reported diagnosis of I/HPAH but clinical features indicative of a different diagnosis. Red-flagged features were the following:
        i. Not meeting hemodynamic diagnostic criteria.
        ii. History of pulmonary embolism or CT scan report suggestive of CTEPH.
        iii. History of structural heart abnormality (PFO/ASD, VSD, PDA) or CT scan/ heart MRI scan indicative of such abnormality.
        iv. Positive autoimmune screen and/or history of connective tissue disease (CTD).
        v. History of left heart disease.
        vi. History of chronic obstructive pulmonary disease.
        vii. History of anorexigen use.
        viii. History of lung fibrosis or CT scan with significant fibrotic changes.
        ix. History of sarcoidosis.

Excel spreadsheets were sent to the centres to verify the diagnosis. Patients identified as non-I/HPAH/PVOD/PCH were suspended and the correct diagnosis was recorded. In all but three individuals diagnostic labels were assigned. Patients with missing diagnostic labels were excluded from the analysis.

#### Missing data assessment

The initial data set was analyzed for missing rate and pattern. Data was found to have a multivariate (many variables had missingness) missing pattern. The transfer coefficient for carbon monoxide (KCO [%pred.]) measured at diagnosis was available for 644 patients

(57%). The missing pattern in KCO in relation to missingness in diagnosis, age at diagnosis and other lung function tests is shown in Figure I in the Data Supplement. Patients with missing KCO data were younger, had better functional status, were more likely non-smokers and had less emphysematous changes on HRCT. Higher missingness was seen more often among prevalent cases (which was caused by the unavailability of old paper medical records), centres that supplied the highest proportion of legacy samples (Imperial and Hammersmith Hospital and Royal Papworth Hospital) and shared care centres (Royal Brompton Hospital). Differences in clinical characteristics between patients with missing and present KCO results are summarized in Table VII.

#### CT scans analysis

The scans available for the repeated analysis were anonymized and transferred to the University of Sheffield, Sheffield, UK. Those CT scans were obtained between 2002 and 2018 (n=269, CT pulmonary angiogram (CTPA) n=241, high resolution computed tomography no CTPA, n=28). Slice thickness was less than 5mm for all studies, typically ≤1mm. Images were analyzed on the open-source software Horos (Annapolis, MD USA). Cardiac and vascular measurements were taken by one observer (MC) and reviewed by the Consultant Radiologist (AS). Thoracic Radiological features were scored semi-quantitatively by two independent Cardiothoracic Radiologist observers each with nine years experience in pulmonary hypertension imaging (AS, SR). To measure the magnitude of agreement between CT scan readers, 22 randomly selected tests were assessed by both radiologists. For categorical variables, the weighted (ordinal data) and unweighted (for non-ordinal data) Cohen's Kappa for two readers were calculated and for continuous variables, the intraclass correlation coefficient (ICC) was computed with the R package "irr" (see Table IV).

#### Selection of phenotypes ('tags') for case-control analysis

Our choice of continuous variables by which we grouped patients was not incidental. Recent registries have shown a considerable shift in PAH demographics[2,3]. Particularly in the ageing western populations, PAH is now diagnosed in elderly patients, with a significant burden of comorbidities, a weaker response to treatment and poorer survival[4]. Although genetic disorders tend to present earlier in life, better phenotypic and genetic characterization of elderly patients is required as this group now constitutes the majority of the adult PAH population. On the other hand, reduced DLCO, which reflects the impairment of alveolar-capillary membrane function, has already been noticed in the analysis of early PAH registry data[5]. Reduced DLCO proved to be an independent predictor of survival, although it does not appear to influence response to treatment with PAH specific therapies[6,7]. Decreased DLCO has also been found in patients with PVOD/PCH with or without biallelic *EIF2AK4* mutations[8–10]. Although some reduction in DLCO is one of the typical features of pulmonary hypertension, PVOD patients show the lowest DLCO values when compared to IPAH or CTEPH. DLNO/DLCO ratio in these patients is highest in PVOD, indicative of proportionally greater involvement of the pulmonary capillary compartment in the disease process[11]. In contrast to these findings, DLCO is relatively preserved in *BMPR2* mutation carriers[12] and shows intermediate values in *BMPR2* mutation carriers with coexisting *EIF2AK4* heterozygous mutations[13]. Contrary to previous reports we collected KCO (the primary measurement) rather than DLCO (the gas exchange potential of the lung at rest and at full inflation). Strong association with survival and a link with other causative mutations makes this phenotype particularly attractive for genetic studies. In summary, we used phenotypes derived from diagnostic classification along with age and KCO strata, which are relevant from a clinical perspective (impact on outcomes and response to treatment) and have previouslybeen associationed with other PAH risk genes.

#### Whole-genome sequencing (WGS), short read alignment, and variant calling and rare variant extraction.

Samples were received as either DNA extracted from whole blood or as whole blood EDTA samples that were extracted at the central DNA extraction and QC laboratory in Cambridge (UK). Following testing for adequate DNA concentration, DNA degradation, and purity, next-generation paired-end whole-genome sequencing was performed on cases and controls using Illumina HiSeq2500 and HiSeq X (Illumina Inc, San Diego, USA) generating three batches with different read lengths: 100bp (377 samples), 125bp (3,154 samples) and 150bp (9,656 samples).

Reads were aligned against the Genome Reference Consortium human genome build 37 (GRCh37, https://www.ncbi.nlm.nih.gov/assembly/GCF_000001405.13/) using the Illumina Isaac Aligner version SAAC00776.15.01.27[14] and variants were called using the Illumina Starling software version 2.1.4.2 (https://support.illumina.com/help/BS_App_TS_Amplicon_OLH_15055858/Content/Source/Informatics/Apps/IsaacVariantCaller_appENR.htm). The variants were then left-aligned, normalized with *bcftools* and loaded into our Hbase database to produce multi-sample variant calls to undertake the genetic association studies[15].

#### Number of PAH domain samples in the analysis

Table I summarizes the number of unrelated samples (estimated using the R/Bioconductor package Genesis[16] and referred to as "maximal unrelated set") by study domain[15]. The PAH domain comprised 1,148 (1,123 unrelated) individuals. Of these, 1,122 cases (following exclusion of three cases with unknown diagnosis and 23 unaffected relatives, Figure 1B) were available for the analysis. A detailed breakdown is provided in Table V. Based on the estimated family networks, only one affected member of each family with the given tag (selected from either the maximal unrelated or the related set) is considered for the genotype-phenotype association testing with BeviMed, which results in a variable number of cases per tag depending on tag availability and family structure. A breakdown of the number of individuals by tag is presented in Table 1. For example, in the case of the PH tag, 1,112 of the 1,122 cases were included in the association testing after excluding nine affected relatives and one case without the tag. The clinical characterization of the entire study population is presented in Table IV in the Data Supplement.

#### "Labelling" of explained cases

For this analysis we made the assumption (widely supported by the literature) that PAH is inherited in Mendelian fashion, meaning that the disease is driven by a single gene. As a consequence of this assumption patients with rare deleterious variants in previously established PAH disease genes (*BMPR2*, *ACVRL1*, *ENG*, *CAV1*, *SMAD1*, *SMAD9*, *KCNK3*, *EIF2AK4*, *TBX4*, *AQP1*, *ATP13A3*, *GDF2*, *SOX17*) that were deemed disease-causing by a genetic multidisciplinary team according to the ACMG Standards and Guidelines[17], were excluded from the association testing for other genes. For example, a patient with a rare deleterious variant in *BMPR2*, which was assessed as pathogenic explaining the phenotype, was not included in the association testing for other genes. Of note, we also reported the results of association testing between rare variants in known PAH risk genes and phenotypes.

#### Selection of gene biotypes and rare variants for testing with BeviMed

The following biotypes of genes defined by Ensembl (https://www.ensembl.org/info/genome/genebuild/biotypes.html) were used for the analysis:

- **lincRNA (long intergenic ncRNA):** Transcripts that are long intergenic non-coding RNA locus with a length >200bp. Requires lack of coding potential and may not be conserved between species.
- **miRNA:** A small RNA (~22bp) that silences the expression of target mRNA.
- **miscRNA:** Miscellaneous RNA. A non-coding RNA that cannot be classified
- **Protein coding:** Gene/transcript that contains an open reading frame (ORF).
- **rRNA:** The RNA component of a ribosome
- **snoRNA:** Small RNA molecules that are found in the cell nucleolus and are involved in the post-transcriptional modification of other RNAs
- **snRNA:** Small RNA molecules that are found in the cell nucleus and are involved in the processing of pre-messenger RNAs
- **TR gene:** T cell receptor gene that undergoes somatic recombination, annotated in collaboration with IMGT http://www.imgt.org/.
    - **TR C gene:** Constant chain T cell receptor gene that undergoes somatic recombination before transcription
    - **TR D gene:** Diversity chain T cell receptor gene that undergoes somatic recombination before transcription
    - **TR J gene:** Joining chain T cell receptor gene that undergoes somatic recombination before transcription
    - **TR V gene:** Variable chain T cell receptor gene that undergoes somatic recombination before transcription
- **IG gene:** Immunoglobulin gene that undergoes somatic recombination, annotated in collaboration with IMGT http://www.imgt.org/.
    - **IG C gene:** Constant chain immunoglobulin gene that undergoes somatic recombination before transcription
    - **IG D gene:** Diversity chain immunoglobulin gene that undergoes somatic recombination before transcription
    - **IG J gene:** Joining chain immunoglobulin gene that undergoes somatic recombination before transcription
    - **IG V gene:** Variable chain immunoglobulin gene that undergoes somatic recombination before transcription
- **Mt_rRNA:** mitochondrial ribosomal RNAs
- **Mt_tRNA:** mitochondrial t-RNAs

Rare variants were extracted from each gene as previously described[15] including a $PMAF_x$ (for a given variant, the probability that the minor allele count is at least the observed minor allele count, given that MAF=1/X) <0.05 with x=1,000 for the recessive and x=10,000 for the dominant association model, and a CADD Phred score ≥10.

To increase power in scenarios where only variants of particular consequence types were associated with the disease risk, association models were fitted to different subsets of variants according to the consequences provided by Ensembl (https://www.ensembl.org/info/genome/variation/prediction/predicted_data.html): the High category, comprised only variants of "high" impact, including likely loss-of-function variants and large deletions; the Moderate category contains variants of impact "moderate", including missense variants or consequence "non_coding_transcript_exon_variant"; the combined category Moderate and High, combining the respective consequence types.

#### Summary of BeviMed methodology

BeviMed is a method for testing association between rare Mendelian disease and a genomic locus by comparing support for a model where disease risks depend on genotypes at rare variant sites in the locus and a genotype-independent 'null' model. The prior probability in such models can vary across variants (reflective of external biological information, i.e. depending on MAF, conservation scores, gene ontologies, expression in the tissue of interest) or be constant for all genes/variants reflecting the prior belief of the overall proportion of variants that are associated with a given phenotype. In the current analysis, we have chosen a constant prior of 0.001 to reflect our assumption that there are around 30 of 32,606 genes that are associated with PAH. We have also calculated the posterior probability for a prior of 0.01 (estimating that 300 genes might be involved in the pathogenesis of PH) and for a prior of 0.00033 (assuming that the total of only ten genes is implicated in the disease) to test the impact of the prior on posterior probability. The results of these analyses are shown in Table XIII. Posterior probability (PP) is a metric that integrates the evidence from observed data (quantified in BF) with the subjectively chosen prior probability ($\pi$) that a gene is truly associated with the phenotype. Because $\pi$ in the genomic analysis is commonly small (ditto), a large BF is required to provide convincing evidence of genotype-phenotype association (PP close to 1). The need for a large BF reflects, therefore, the small number of expected true genotype-phenotype associations and is comparable to FDR adjustment in frequentist approaches with the difference that frequentists adjust for the number of comparisons. Rather than relying on significance thresholds (used in the frequentist framework but discouraged in the Bayesian approach) we ranked our findings according to the logarithm of BF/probability (in case of constant priors the ranking for both parameters is the same) truncating the list around the level last previously reported PAH risk gene for the prior set at 0.001 which we believe is best justified in this case.

#### Descriptive statistics

Statistical analysis and data visualization were performed in R (www.r-project.org). Summary statistics are shown as mean (±SD) or median [IQR] according to data distribution. The number of available data points is reported in tables. Comparisons between the categorical variables were performed using Fisher's exact and Chi-square test, comparisons between continuous non-normally distributed variables were performed with the Mann-Whitney test or the Kruskal-Wallis test. Adjustment for multiple comparisons was performed when appropriate. The Kaplan-Meier method was used to visualize survival curves; the log-rank test was used to compare survival between two or more groups; Cox proportional hazards regression was used to examine the effect of variables on survival. Testing for the proportional hazards assumption was done, and the assumptions were met.

### Results

#### Summary of identified variants in previously reported genes

We identified single nucleotide variants (SNVs), small insertions and deletions (indels) and larger deletions (see Figure IV) in previously established genes (namely, *BMPR2*, *ACVRL1*, *ENG*, *SMAD1*, *SMAD4*, *SMAD9*, *KCNK3*, *TBX4*, *EIF2AK4*, *AQP1*, *ATP13A3*, *GDF2*, *SOX17*) in 271 (24.2%) of the 1,122 cases recruited to the PAH domain and interpreted them based on the ACMG standards and guidelines[17]. We identified 124 (45.8%) and 110 (40.6%) subjects carrying pathogenic and likely pathogenic variants, respectively, and 37 individuals (13.6%) with variants of uncertain significance. Of these, 161 (59.4% of explained, 14.3% of all cases) carried pathogenic mutations in *BMPR2*, 96 subjects (59.6%) carried loss-of-function variants (including stop gained, frameshift and splice donor/acceptor variants), 27 (16.8%) possessed larger deletions and 38 cases (23.6%) presented with deleterious, missense variants. Twenty-one cases had rare variants in *TBX4* (7.7% of explained, 1.9% of all cases) comprising of either loss-of-function mutations (10 subjects (3.7%)), larger deletions (two cases (0.7%)) or deleterious missense variants (nine individuals (3.3%)). Mutations in *ATP13A3* (15 cases; 5.5% of explained and 1.3% of all cases) comprised loss-of-function mutations (six cases (2.2%)) and missense mutations (nine cases (3.3%), including one homozygous mutation in a child already reported in Barozzi *et al.*[18]). We identified 14 patients (5.2% of explained, 1.2% of all cases) with biallelic *EIF2AK4* mutations, six homozygous (2.2%) and eight compound heterozygous (3%) individuals. A further 14 cases (5.2% of explained, 1.2% of all cases) carried mutations in *GDF2*, the majority exhibited deleterious missense mutations (10 individuals (3.7%)), two cases (0.7%) harboured large deletions and another two cases (0.7%) had loss-of-function variants which co-occurred with a large deletion in *BMPR2* and a heterozygous missense variant in *SMAD9* respectively. *ACVRL1* mutations were detected in 11 cases (4.1% of explained, 1% of all cases), with the majority being heterozygous missense variants (10 cases (3.7%)). Only one individual (0.4%) had a loss-of-function variant *ACVRL1*. Heterozygous missense variants in *AQP1* were revealed in nine subjects (3.3% of explained, 0.8% of all cases), of which one was accompanied with a heterozygous missense variant in *BMPR2*. An additional case presented with a loss-of-function variant in *AQP1*. In *ENG* we predominantly identified heterozygous missense variants in eight cases (3% of explained, 0.7% of all cases), of which two co-occurred with a large deletion in *BMPR2* each, and a loss-of-function variant in one case (0.4% of explained, 0.1% of all cases). Another nine cases (3.3% of explained, 0.8% of all cases) harboured variants in *SOX17*, three with loss-of-function and six with heterozygous missense variants. *SMAD9* variants were found in eight cases (3% of explained, 0.2% of all cases), six had heterozygous missense variants, of which two co-occurred with loss-of-function variants in *BMPR2* and *GDF2*, and two possessed heterozygous missense mutations, of which one was accompanied by a heterozygous missense variant. A further four patients (1.5% of explained, 0.4% of all cases) exhibited deleterious missense (three cases (1.1%)) and loss-of-function (one case (0.4%)) variants in *KCNK3*. Finally, two cases (0.7% of explained, 0.2% of all cases) harboured a loss-of-function variant in *SMAD1*, of which one co-existed with a loss-of-function variant in *BMPR2*.

### Supplemental references

1. Pang C, Sollie A, Sijtsma A, Hendriksen D, Charbon B, de Haan M, de Boer T, Kelpin F, Jetten J, van der Velde JK, Smidt N, Sijmons R, Hillege H, Swertz MA. SORTA: a system for ontology-based re-coding and technical annotation of biomedical phenotype data

[Internet]. Database. 2015;2015:bav089. Available from: http://dx.doi.org/10.1093/database/bav089

2. Test VJ, Farber HW, McGoon MD, Parsons L, Channick RN. Pulmonary Arterial Hypertension in the Elderly: Baseline Characteristics and Evaluation of Therapeutics. An Examination of the Reveal Registry [Internet]. B27. FROM ALPHA TO OMEGA: ASSESSMENT AND OUTCOMES IN PULMONARY HYPERTENSION. 2009;Available from: http://dx.doi.org/10.1164/ajrccm-conference.2009.179.1_meetingabstracts.a2649

3. Ling Y, Johnson MK, Kiely DG, Condliffe R, Elliot CA, Gibbs JSR, Howard LS, Pepke-Zaba J, Sheares KKK, Corris PA, Fisher AJ, Lordan JL, Gaine S, Coghlan JG, Wort SJ, Gatzoulis MA, Peacock AJ. Changing demographics, epidemiology, and survival of incident pulmonary arterial hypertension: results from the pulmonary hypertension registry of the United Kingdom and Ireland. *Am J Respir Crit Care Med*. 2012;186:790–796.

4. Hoeper MM, Huscher D, Ghofrani HA, Delcroix M, Distler O, Schweiger C, Grunig E, Staehler G, Rosenkranz S, Halank M, Held M, Grohé C, Lange TJ, Behr J, Klose H, Wilkens H, Filusch A, Germann M, Ewert R, Seyfarth H-J, Olsson KM, Opitz CF, Gaine SP, Vizza CD, Vonk-Noordegraaf A, Kaemmerer H, Gibbs JSR, Pittrow D. Elderly patients diagnosed with idiopathic pulmonary arterial hypertension: results from the COMPERA registry. *Int J Cardiol*. 2013;168:871–880.

5. Test VJ, Farber HW, McGoon MD, Parsons L, Channick RN. Pulmonary Arterial Hypertension in the Elderly: Baseline Characteristics and Evaluation of Therapeutics. An Examination of the Reveal Registry [Internet]. B27. FROM ALPHA TO OMEGA: ASSESSMENT AND OUTCOMES IN PULMONARY HYPERTENSION. 2009;Available from: http://dx.doi.org/10.1164/ajrccm-conference.2009.179.1_meetingabstracts.a2649

6. Trip P, Girerd B, Bogaard H-J, de Man FS, Boonstra A, Garcia G, Humbert M, Montani

D, Vonk-Noordegraaf A. Diffusion capacity and BMPR2 mutations in pulmonary arterial hypertension. *Eur Respir J*. 2014;43:1195–1198.

15. Turro E, Astle WJ, Megy K, Gräf S, Greene D, Shamardina O, Allen HL, Sanchis-Juan A, Frontini M, Thys C, Stephens J, Mapeta R, Burren OS, Downes K, Haimel M, Tuna S, Deevi SVV, Aitman TJ, Bennett DL, Calleja P, Carss K, Caulfield MJ, Chinnery PF, Dixon PH, Gale DP, James R, Koziell A, Laffan MA, Levine AP, Maher ER, Markus HS, Morales J, Morrell NW, Mumford AD, Ormondroyd E, Rankin S, Rendon A, Richardson S, Roberts I, Roy NBA, Saleem MA, Smith KGC, Stark H, Tan RYY, Themistocleous AC, Thrasher AJ, Watkins H, Webster AR, Wilkins MR, Williamson C, Whitworth J, Humphray S, Bentley DR, NIHR BioResource for the 100,000 Genomes Project, Kingston N, Walker N, Bradley JR, Ashford S, Penkett CJ, Freson K, Stirrups KE, Raymond FL, Ouwehand WH. Whole-genome sequencing of patients with rare diseases in a national health system. *Nature*. 2020;583:96–102.

16. Gogarten SM, Sofer T, Chen H, Yu C, Brody JA, Thornton TA, Rice KM, Conomos MP. Genetic association testing using the GENESIS R/Bioconductor package [Internet]. Bioinformatics. 2019;Available from: http://dx.doi.org/10.1093/bioinformatics/btz567

17. Richards S, Aziz N, Bale S, Bick D, Das S, Gastier-Foster J, Grody WW, Hegde M, Lyon E, Spector E, Voelkerding K, Rehm HL, ACMG Laboratory Quality Assurance Committee. Standards and guidelines for the interpretation of sequence variants: a joint consensus recommendation of the American College of Medical Genetics and Genomics and the Association for Molecular Pathology. *Genet Med*. 2015;17:405–424.

18. Barozzi C, Galletti M, Tomasi L, De Fanti S, Palazzini M, Manes A, Sazzini M, Galiè N. A Combined Targeted and Whole Exome Sequencing Approach Identified Novel Candidate Genes Involved in Heritable Pulmonary Arterial Hypertension. *Sci Rep*. 2019;9:753.

#### Supplemental figure legends

<u>Figure I</u>. **Summary of missing data. A**, The fraction of missing data for KCO, in comparison to diagnosis, age at diagnosis and lung function tests. **B**, The missingness pattern in KCO, in relation to diagnosis, age at diagnosis and lung function tests. KCO - transfer coefficient for carbon monoxide. $FEV_1$ - forced expiratory volume in 1 second, FVC - forced vital capacity, TLC - total lung capacity.

<u>Figure II</u>. **Flowchart describing the definition of diagnostic and phenotypic tags.** A detailed description is provided in the supplemental material. The definition of tags is listed in <u>Table 1</u>.

<u>Figure III</u>. **Characterization and survival analysis of the cohort based on transfer coefficient for carbon monoxide (KCO) and age at diagnosis.** Distribution of and Kaplan-Meier survival curves for KCO below and above the 50% predicted threshold (A, B), KCO tertiles (C D), and age tertiles (E, F).

<u>Figure IV</u>. **Summary of large deletions identified in previously established disease genes.** Deletions are indicated by light blue boxes. The protein-coding genes, annotated in the displayed region by Ensembl (GRCh37, version 75), are depicted in the bottom panels. The affected genomic regions, with the disease gene locus highlighted in red and the magnified view focusing on the gene loci, are shown for *BMPR2* (**A**, **B**), *GDF2* (**C**, **D**) and *TBX4* (**E**, **F**).

<u>Figure V</u>. **Chest computerized tomography (CT) scans of the daughter of W00229 carrying the inherited nonsense variant in *KDR*. A**, The scan shows bibasal reticular ground glass changes with mild traction bronchiectasis. **B**, Upper left lobe shows subpleural

reticular ground glass changes in keeping with interstitial fibrosis. Further ground glass changes are visible in the right lung.

#### Supplemental table legends

**Table I**. NIHR BioResource - Rare Diseases domain definitions and number of unrelated individuals per project.

**Table II.** Summary of electronic case report forms constructed to capture phenotypic information.

**Table III.** Reporting proforma for CT scan revision. Abbreviations: CTPA - Computerized Tomography Pulmonary Angiogram, HRCT - High-Resolution Computerized Tomography, GGO - ground-glass opacities.

**Table IV.** Summary of imaging analysis. Intra-rater reliability: GGO centrilobular pattern severity weighted Cohen's Kappa=0.679, p-value <0.001; GGO distribution unweighted Cohen's Kappa = 1, p-value 0.046; Severity of GGO non-specific pattern - no positive findings; Pulmonary arteriovenous malformations - no positive findings; largest BRA size - no positive findings; Mediastinal venous collaterals: unweighted Cohen's Kappa = 1, p-value <0.001; Intralobular septal thickening weighted Cohen's Kappa = 1, p-value <0.001; Mediastinal lymphadenopathy unweighted Cohen's Kappa=0.83, p-value <0.001; Mediastinal lymphadenopathy size [mm] intraclass correlation coefficient (ICC) 0.717, p-value 0.088; Emphysema - not enough positive findings, Bronchial wall thickening - not enough positive findings, Fibrosis - no positive findings; Pleural effusion weighted Cohen's Kappa 0.826, p-value <0.001; Air trapping weighted Cohen's Kappa 0.845, p-value <0.001; Subpleural scarring - not enough positive findings. Abbreviations: IPAH - idiopathic pulmonary arterial hypertension, HPAH - hereditary pulmonary arterial hypertension, PVOD - pulmonary veno-occlusive disease, PCH - Pulmonary capillary haemangiomatosis, GGO - ground-glass opacities, BA - bronchial artery, C - central, U - upper, Z - zonal, D - diffuse, bial. - biallelic, mis. - missense, lof - loss-of-function.

**Table V.** Number of PAH domain samples in the analysis broken down by maximal unrelated set and related set, and cases (included and excluded) and unaffected relatives.

**Table VI.** Clinical characterization of the study population. Entire cohort (n=1,122) was composed of IPAH (n=972), HPAH (n=73), PVOD/PCH (n=20), PH associated with left heart disease (n=7), PH associated with lung disease (n=8), chronic thromboembolic pulmonary hypertension (n=6), multifactorial PH (n=6), hereditary hemorrhagic telangiectasia (n=1). Abbreviations: BMI - body mass index, WHO FC - World Health Organization Functional Class, 6MWD - 6-minute walk distance, mRAP - mean right atrial pressure, mPAP - mean pulmonary artery pressure, CO - cardiac output, $FEV_1$ - forced expiratory capacity in 1 second, FVC - forced vital capacity, KCO - transfer factor coefficient for carbon monoxide, Hb - hemoglobin, RDW - red cell distribution width, WBC - white blood cell count, NTproBNP - N-terminal pro B-type natriuretic peptide, BNP - B-type natriuretic peptide, CRP - C-Reactive Protein Protein, HTN - hypertension, DM - diabetes mellitus, CAD - coronary artery disease, CVA - cerebrovascular accident, COPD - chronic obstructive pulmonary disease, CCB - calcium channel blocker, ERA - endothelin receptor antagonists, PA - prostacyclin analogs, PED5 - phosphodiesterase type 5, sGC - soluble guanylate cyclase.

**Table VII.** Clinical differences between patients with present and absent KCO. Abbreviations: BMI - body mass index, WHO FC - World Health Organization Functional Class, 6MWD - 6-minute walk distance, mRAP - mean right atrial pressure, mPAP - mean pulmonary artery pressure, PAWP - Pulmonary Artery Wedge Pressure, CI - cardiac index, PVR - pulmonary vascular resistance, $SvO_2$ [%] - mixed venous saturation, HRCT - High-Resolution Computerized Tomography, $FEV_1$ - forced expiratory capacity in 1 second, FVC - forced vital capacity, COPD - chronic obstructive pulmonary disease, OSA - obstructive sleep apnoea.

**Table VIII.** Clinical characteristics study individuals by KCO threshold. None of the patients had systemic lupus erythematosus, systemic sclerosis, undifferentiated connective tissue disease or ankylosing spondylitis. Abbreviations: BMI - body mass index, WHO FC - World Health Organization Functional Class, 6MWD - 6-minute walk distance, $SpO_2$ - peripheral capillary oxygen saturation, mRAP - mean right atrial pressure, mPAP - mean pulmonary artery pressure, PAWP - Pulmonary Artery Wedge Pressure, CO - cardiac output, $SvO_2$ - Mixed venous oxygen saturation, $FEV_1$ - forced expiratory capacity in 1 second, FVC - forced vital capacity, TLC - Total Lung Capacity, KCO - transfer coefficient for carbon monoxide, HRCT - High-Resolution Computerized Tomography, NTproBNP - N-terminal pro B-Type Natriuretic Peptide, BNP - B-Type Natriuretic Peptide, CRP - C-Reactive Protein Protein, Hb - hemoglobin, WBC - white blood cell count, COPD - chronic obstructive pulmonary disease, OSA - obstructive sleep apnoea, CAD - coronary artery disease, CVA - cerebrovascular accident, PAD - peripheral artery disease, HTN - hypertension, DM - diabetes mellitus.

**Table IX.** Clinical characteristics of study individuals by KCO tertiles. None of the patients had systemic lupus erythematosus, systemic sclerosis, undifferentiated connective tissue disease or ankylosing spondylitis. Abbreviations: BMI - body mass index, WHO FC - World Health Organization Functional Class, 6MWD - 6-minute walk distance, $SpO_2$ - peripheral capillary oxygen saturation, mRAP - mean right atrial pressure, mPAP - mean pulmonary artery pressure, PAWP - Pulmonary Artery Wedge Pressure, CO - cardiac output, $SvO_2$ - mixed venous oxygen saturation, NO - nitric oxide, $FEV_1$ - forced expiratory capacity in 1 second, FVC - forced vital capacity, TLC - Total Lung Capacity, KCO - transfer coefficient for carbon monoxide, HRCT - High-Resolution Computerized Tomography, NTproBNP - N-terminal pro B-type natriuretic peptide, BNP - B-type natriuretic peptide, CRP - C-Reactive Protein Protein, Hb - hemoglobin, WBC - white blood cell count, COPD - chronic obstructive pulmonary disease, OSA - obstructive sleep apnoea, CAD - coronary artery disease, CVA - cerebrovascular accident, PAD - peripheral artery disease, HTN - hypertension, DM - diabetes mellitus.

**Table X.** Results of Cox regression analysis relating overall survival to selected variables at baseline. Abbreviations: CI - Confidence interval, 6MWD - 6-minute walking distance, mPAP - mean pulmonary arterial pressure, mRAP - mean right atrial pressure, PVR - pulmonary vascular resistance, WU - Wood units, KCO - transfer coefficient for carbon monoxide, CAD - coronary artery disease, COPD - chronic obstructive pulmonary disease, HTN - systemic hypertension, HRCT - High-Resolution Computerized Tomography.

**Table XI.** Clinical characteristics of all individuals by age tertiles. None of the patients had undifferentiated connective tissue disease, incident cases were defined as those diagnosed within 6 months from study commencement. Abbreviations: BMI - body mass index, WHO FC - World Health Organization Functional Class, 6MWD - 6-minute walk distance, $SpO_2$ - peripheral capillary oxygen saturation, mRAP - mean right atrial pressure, mPAP - mean pulmonary artery pressure, PAWP - Pulmonary Artery Wedge Pressure, CO - cardiac output, $SvO_2$ - Mixed venous oxygen saturation, NO - nitric oxide, $FEV_1$ - forced expiratory capacity in 1 second, FVC - forced vital capacity, TLC - Total Lung Capacity, KCO - transfer factor coefficient for carbon monoxide, NTproBNP - N-terminal pro B-type natriuretic peptide, BNP - B-type natriuretic peptide, CRP - C-Reactive Protein Protein, Hb - hemoglobin, WBC - white blood cell count, COPD - chronic obstructive pulmonary disease, OSA - obstructive sleep apnoea, CAD - coronary artery disease, CVA - cerebrovascular accident, PAD - peripheral artery disease, HTN - hypertension, DM - diabetes mellitus.

**Table XII.** Clinical characteristics of IPAH patients who harbor loss-of-function variants in *BMPR2*, *EIF2AK4*, and *KDR*. Comorbidities are reported as the number and percentage of cases possessing a disease entity. Abbreviations: BMI - body mass index, WHO FC - World Health Organization functional class, 6MWD - 6-minute walk distance, $SpO_2$ - arterial oxygen saturation, mRAP - mean right atrial pressure, mPAP - mean pulmonary artery pressure, mPAWP - mean pulmonary artery wedge pressure, CO - cardiac output, PVR - pulmonary vascular resistance, NO - nitric oxide challenge, $FEV_1$ - forced expiratory volume in 1 second, FVC - forced vital capacity, KCO - transfer factor coefficient for carbon monoxide, COPD - chronic obstructive pulmonary disease, OSA - obstructive sleep apnea, CAD - coronary artery disease, HTN - systemic hypertension, CKD - chronic kidney disease, Hb - hemoglobin, WBC - white blood cells, TSH - thyroid-stimulating hormone, bial. - biallelic, lof - loss-of-function.

**Table XIII.** BeviMed analysis result comparison. Shown are Bayes factors (BF) and posterior probabilities (PP) for three different priors (**π**).

### Supplemental Tables

**Table I.** NIHR BioResource - Rare Diseases domain definitions and number of maximal unrelated individuals per project.

| Project acronym | Project | Number of individuals |
|---|---|---|
| GEL | Genomics England Ltd | 3058 |
| BPD | Bleeding, Thrombotic and Platelet Disorders | 986 |
| PID | Primary Immune Disorders | 1027 |
| CNTRL | Processed Controls | 50 |
| IRD | Inherited Retinal Disorders | 717 |
| NDD | Neurological and Developmental Disorders | 518 |
| EDS | Ehlers Danlos Syndrome | 15 |
| HCM | Hypertrophic Cardiomyopathy | 239 |
| PMG | Primary Membranoproliferative Glomerulonephritis | 181 |
| SRNS | Steroid Resistant Nephrotic Syndrome | 234 |
| CSVD | Cerebral Small Vessel Disease | 134 |
| NPD | Neuropathic Pain Disorder | 185 |
| ICP | Intrahepatic Cholestasis of Pregnancy | 267 |
| LHON | Leber Hereditary Optic Neuropathy | 54 |
| MPMT | Multiple Primary Tumours | 554 |
| SMD | Stem Cell & Myeloid Disorders | 153 |
| PAH | Pulmonary arterial hypertension | 1123 |
| UKBio | UK BioBank | 764 |
| | Total | 10259 |

**Table II.** Summary of electronic case report forms constructed to capture phenotypic information.

| |
|---|
| ID capture |
| Demographics |
| Functional class |
| Clinical features by history |
| Clinical features by examination |
| Risk factors |
| Haemodynamics |
| Echocardiography |
| Electrocardiogram |
| Lung function |
| Associated Diseases |
| Clinical blood tests |
| Survival |
| Arterial blood gases |
| Imaging |
| Exercise performance |
| Body system |
| Drug treatment history (PAH) |
| Drug treatment history (other) |
| Family history |
| Epidemiology questionnaire |

**Table III.** Reporting proforma for CT scan revision. Abbreviations: CTPA - Computerized Tomography Pulmonary Angiogram, HRCT - High-Resolution Computerized Tomography, GGO - ground-glass opacities.

| Parameter | Response |
|---|---|
| ID | character |
| Reader | character |
| CT scan date | date |
| Slice thickness | numeric |
| Number of slices | numeric |
| CTPA | done/not done |
| HRCT | done/not done |
| Expiratory CT | done/not done |
| Pleural effusion | Nil; Trace; Mild; Moderate; Severe |
| Subcutaneous oedema | present, absent |
| Severity of GGO centrilobular pattern | Nil; Trace; Mild; Moderate; Severe |
| Severity of GGO non-specific mosaic pattern | Nil; Trace; Mild; Moderate; Severe |
| Distribution of GGO | C-central; U-upper; Z-zonal; D-diffuse |
| Pulmonary arteriovenous malformations | present, absent |
| Largest bronchial artery size | numeric [mm] |
| Mediastinal venous collaterals | present, absent |
| Intralobular septal thickening | Nil; Trace; Mild; Moderate; Severe |
| Mediastinal lymphadenopathy | present, absent |
| Mediastinal lymphadenopathy subcarinal | [mm] |
| Emphysema | Nil; Mild; Moderate; Severe |
| Bronchial wall thickening | Nil; Trace; Mild; Moderate; Severe |
| Fibrosis | Nil; Mild; Moderate; Severe |
| Air trapping | Nil; Trace; Mild; Moderate; Severe |
| Subpleural scarring | Nil; Mild; Moderate; Severe |

| **Table IV.** Summary of imaging analysis. Intra-rater reliability: GGO centrilobular pattern severity weighted Cohen's Kappa=0.679, p-value <0.001; GGO distribution unweighted Cohen's Kappa = 1, p-value 0.046; Severity of GGO non-specific pattern - no positive findings; Pulmonary arteriovenous malformations - no positive findings; largest BRA size - no positive findings; Mediastinal venous collaterals: unweighted Cohen's Kappa = 1, p-value <0.001; Intralobular septal thickening weighted Cohen's Kappa = 1, p-value <0.001; Mediastinal lymphadenopathy unweighted Cohen's Kappa=0.83, p-value <0.001; Mediastinal lymphadenopathy size [mm] intraclass correlation coefficient (ICC) 0.717, p-value 0.088; Emphysema - not enough positive findings, Bronchial wall thickening - not enough positive findings, Fibrosis - no positive findings; Pleural effusion weighted Cohen's Kappa 0.826, p-value <0.001; Air trapping weighted Cohen's Kappa 0.845, p-value <0.001; Subpleural scarring - not enough positive findings. Abbreviations: IPAH - idiopathic pulmonary arterial hypertension, HPAH - hereditary pulmonary arterial hypertension, PVOD - pulmonary veno-occlusive disease, PCH - Pulmonary capillary haemangiomatosis, GGO - ground-glass opacities, BA - bronchial artery, C - central, U - upper, Z - zonal, D - diffuse, bial. - biallelic, mis. - missense, lof - loss-of-function. | | | | | | | | | | |
|---|---|---|---|---|---|---|---|---|---|---|
| | **ALL (N=267)** | ***BMPR2* (N=44)** | ***EIF2AK4* (N=6)** | ***EIF2AK4* (bial.) (N=7)** | ***KDR* (mis.) (N=5)** | ***KDR* (lof) N=4** | **No mutation (N=183)** | **Other mutations (N=18)** | **p overall** | **N** |
| **Sex: female** | 177 (66.3%) | 28 (63.6%) | 5 (83.3%) | 3 (42.9%) | 3 (60.0%) | 2 (50.0%) | 127 (69.4%) | 9 (50.0%) | 0.364 | 267 |
| **Age at diagnosis [years]** | 50.1 (17.0) | 44.2 (13.7) | 51.0 (16.7) | 28.5 (10.8) | 36.6 (13.1) | 65.2 (4.3) | 52.9 (17.2) | 44.4 (14.1) | <0.001 | 266 |
| **Diagnosis verified:** | | | | | | | | | 0.009 | 267 |
| **HPAH** | 20 (7.5%) | 13 (29.5%) | 0 (0.0%) | 0 (0.0%) | 0 (0.0%) | 0 (0.0%) | 4 (2.2%) | 3 (16.7%) | | |
| **IPAH** | 235 (88.0%) | 31 (70.5%) | 5 (83.3%) | 4 (57.1%) | 5 (100.0%) | 4 (100.0%) | 171 (93.4%) | 15 (83.3%) | | |
| **PCH** | 1 (0.4%) | 0 (0.0%) | 0 (0.0%) | 1 (14.3%) | 0 (0.0%) | 0 (0.0%) | 0 (0.0%) | 0 (0.0%) | | |
| **PVOD** | 11 (4.1%) | 0 (0.0%) | 1 (16.7%) | 2 (28.6%) | 0 (0.0%) | 0 (0.0%) | 8 (4.4%) | 0 (0.0%) | | |
| **GGO centrilobular pattern severity:** | | | | | | | | | 0.047 | |
| **Nil** | 22 (50.0) | 4 (66.7) | 2 (28.6) | 4 (80.0) | 2 (50.0) | 119 (65.0) | 12 (66.7) | 165 (61.8) | | |
| **Trace** | 3 (6.8) | 0 (0.0) | 1 (14.3) | 0 (0.0) | 0 (0.0) | 23 (12.6) | 2 (11.1) | 29 (10.9) | | |
| **Mild** | 9 (20.5) | 0 (0.0) | 1 (14.3) | 0 (0.0) | 2 (50.0) | 16 (8.7) | 1 (5.6) | 29 (10.9) | | |
| **Moderate** | 3 (6.8) | 0 (0.0) | 1 (14.3) | 1 (20.0) | 0 (0.0) | 17 (9.3) | 2 (11.1) | 24 (9.0) | | |
| **Severe** | 7 (15.9) | 2 (33.3) | 2 (28.6) | 0 (0.0) | 0 (0.0) | 8 (4.4) | 1 (5.6) | 20 (7.5) | | |
| **Severity of GGO non-specific pattern:** | | | | | | | | | 0.723 | 267 |

| | | | | | | | | | | |
|---|---|---|---|---|---|---|---|---|---|---|
| **Nil** | 238 (89.1%) | 42 (95.5%) | 3 (50.0%) | 5 (71.4%) | 5 (100.0%) | 2 (50.0%) | 165 (90.2%) | 16 (88.9%) | | |
| **Trace** | 10 (3.7%) | 1 (2.3%) | 2 (33.3%) | 0 (0.0%) | 0 (0.0%) | 1 (25.0%) | 6 (3.3%) | 0 (0.0%) | | |
| **Mild** | 11 (4.1%) | 1 (2.3%) | 0 (0.0%) | 1 (14.3%) | 0 (0.0%) | 1 (25.0%) | 7 (3.8%) | 1 (5.6%) | | |
| **Moderate** | 7 (2.6%) | 0 (0.0%) | 1 (16.7%) | 1 (14.3%) | 0 (0.0%) | 0 (0.0%) | 4 (2.2%) | 1 (5.6%) | | |
| **Severe** | 1 (0.4%) | 0 (0.0%) | 0 (0.0%) | 0 (0.0%) | 0 (0.0%) | 0 (0.0%) | 1 (0.5%) | 0 (0.0%) | | |
| **Distribution of GGO:** | | | | | | | | | 0.108 | 122 |
| **C** | 11 (9.0%) | 0 (0.0%) | 1 (20.0%) | 2 (40.0%) | 0 (0.0%) | 2 (66.7%) | 6 (7.7%) | 0 (0.0%) | | |
| **D** | 81 (66.4%) | 19 (90.5%) | 3 (60.0%) | 3 (60.0%) | 2 (100.0%) | 1 (33.3%) | 48 (61.5%) | 5 (62.5%) | | |
| **U** | 14 (11.5%) | 1 (4.8%) | 0 (0.0%) | 0 (0.0%) | 0 (0.0%) | 0 (0.0%) | 12 (15.4%) | 1 (12.5%) | | |
| **Z** | 16 (13.1%) | 1 (4.8%) | 1 (20.0%) | 0 (0.0%) | 0 (0.0%) | 0 (0.0%) | 12 (15.4%) | 2 (25.0%) | | |
| **Pulmonary arteriovenous malformations: Yes** | 4 (1.6%) | 3 (7.3%) | 0 (0.0%) | 0 (0.0%) | 0 (0.0%) | 0 (0.0%) | 1 (0.6%) | 0 (0.0%) | 0.155 | 244 |
| **Largest bronchial artery size** | 3.0 (0.6) | 3.4 (0.5) | 3.0 (.) | 2.5 (0.2) | . (.) | . (.) | 2.5 (0.4) | 4.0 (.) | 0.037 | 12 |
| **Mediastinal venous collaterals: Yes** | 204 (94.0%) | 37 (100.0%) | 1 (100.0%) | 0 (.%) | 5 (100.0%) | 3 (100.0%) | 145 (91.8%) | 13 (100.0%) | 0.383 | 217 |
| **Intralobular septal thickening:** | | | | | | | | | 0.113 | 241 |
| **Nil** | 214 (88.8%) | 37 (92.5%) | 0 (0.0%) | 0 (.%) | 4 (80.0%) | 4 (100.0%) | 153 (88.4%) | 16 (88.9%) | | |
| **Trace** | 15 (6.2%) | 2 (5.0%) | 0 (0.0%) | 0 (.%) | 1 (20.0%) | 0 (0.0%) | 12 (6.9%) | 0 (0.0%) | | |
| **Mild** | 9 (3.7%) | 1 (2.5%) | 0 (0.0%) | 0 (.%) | 0 (0.0%) | 0 (0.0%) | 7 (4.0%) | 1 (5.6%) | | |
| **Moderate** | 3 (1.2%) | 0 (0.0%) | 1 (100.0%) | 0 (.%) | 0 (0.0%) | 0 (0.0%) | 1 (0.6%) | 1 (5.6%) | | |
| **Mediastinal lymphoaenopathy: Yes** | 216 (81.2%) | 40 (90.9%) | 3 (50.0%) | 3 (42.9%) | 2 (50.0%) | 1 (25.0%) | 149 (81.4%) | 18 (100.0%) | <0.001 | 266 |
| **Mediastinal lymphoadenopathy size [mm]** | 14.9 (4.0) | 12.5 (1.3) | 17.0 (3.6) | 14.8 (2.4) | 15.5 (0.7) | 11.0 (0.0) | 15.3 (4.4) | . (.) | 0.354 | 50 |

| | | | | | | | | | | |
|---|---|---|---|---|---|---|---|---|---|---|
| **Emphysema:** | | | | | | | | | 0.819 | 267 |
| **Nil** | 233 (87.3%) | 38 (86.4%) | 5 (83.3%) | 7 (100.0%) | 4 (80.0%) | 3 (75.0%) | 159 (86.9%) | 17 (94.4%) | | |
| **Trace** | 20 (7.5%) | 4 (9.1%) | 1 (16.7%) | 0 (0.0%) | 1 (20.0%) | 0 (0.0%) | 13 (7.1%) | 1 (5.6%) | | |
| **Mild** | 8 (3.0%) | 1 (2.3%) | 0 (0.0%) | 0 (0.0%) | 0 (0.0%) | 1 (25.0%) | 6 (3.3%) | 0 (0.0%) | | |
| **Moderate** | 6 (2.2%) | 1 (2.3%) | 0 (0.0%) | 0 (0.0%) | 0 (0.0%) | 0 (0.0%) | 5 (2.7%) | 0 (0.0%) | | |
| **Bronchial wall thickening:** | | | | | | | | | 0.049 | 241 |
| **Nil** | 210 (87.1%) | 33 (82.5%) | 0 (0.0%) | 0 (.%) | 5 (100.0%) | 3 (75.0%) | 156 (90.2%) | 13 (72.2%) | | |
| **Trace** | 18 (7.5%) | 5 (12.5%) | 1 (100.0%) | 0 (.%) | 0 (0.0%) | 0 (0.0%) | 10 (5.8%) | 2 (11.1%) | | |
| **Mild** | 11 (4.6%) | 2 (5.0%) | 0 (0.0%) | 0 (.%) | 0 (0.0%) | 1 (25.0%) | 6 (3.5%) | 2 (11.1%) | | |
| **Moderate** | 2 (0.8%) | 0 (0.0%) | 0 (0.0%) | 0 (.%) | 0 (0.0%) | 0 (0.0%) | 1 (0.6%) | 1 (5.6%) | | |
| **Fibrosis:** | | | | | | | | | 0.06 | 267 |
| **Nil** | 255 (95.5%) | 43 (97.7%) | 6 (100.0%) | 7 (100.0%) | 5 (100.0%) | 2 (50.0%) | 175 (95.6%) | 17 (94.4%) | | |
| **Trace** | 5 (1.9%) | 0 (0.0%) | 0 (0.0%) | 0 (0.0%) | 0 (0.0%) | 0 (0.0%) | 5 (2.7%) | 0 (0.0%) | | |
| **Mild** | 6 (2.2%) | 1 (2.3%) | 0 (0.0%) | 0 (0.0%) | 0 (0.0%) | 2 (50.0%) | 3 (1.6%) | 0 (0.0%) | | |
| **Moderate** | 1 (0.4%) | 0 (0.0%) | 0 (0.0%) | 0 (0.0%) | 0 (0.0%) | 0 (0.0%) | 0 (0.0%) | 1 (5.6%) | | |
| **Pleural effusion:** | | | | | | | | | 0.368 | 267 |
| **Nil** | 240 (89.9%) | 41 (93.2%) | 5 (83.3%) | 7 (100.0%) | 4 (80.0%) | 3 (75.0%) | 165 (90.2%) | 15 (83.3%) | | |
| **Trace** | 11 (4.1%) | 1 (2.3%) | 1 (16.7%) | 0 (0.0%) | 0 (0.0%) | 0 (0.0%) | 6 (3.3%) | 3 (16.7%) | | |
| **Mild** | 7 (2.6%) | 2 (4.5%) | 0 (0.0%) | 0 (0.0%) | 1 (20.0%) | 0 (0.0%) | 4 (2.2%) | 0 (0.0%) | | |
| **Moderate** | 7 (2.6%) | 0 (0.0%) | 0 (0.0%) | 0 (0.0%) | 0 (0.0%) | 1 (25.0%) | 6 (3.3%) | 0 (0.0%) | | |
| **Severe** | 2 (0.7%) | 0 (0.0%) | 0 (0.0%) | 0 (0.0%) | 0 (0.0%) | 0 (0.0%) | 2 (1.1%) | 0 (0.0%) | | |

| | | | | | | | | | | |
|---|---|---|---|---|---|---|---|---|---|---|
| **Air trapping:** | | | | | | | | | 0.085 | 267 |
| **Nil** | 214 (80.1%) | 40 (90.9%) | 5 (83.3%) | 7 (100.0%) | 4 (80.0%) | 1 (25.0%) | 144 (78.7%) | 13 (72.2%) | | |
| **Trace** | 26 (9.7%) | 3 (6.8%) | 1 (16.7%) | 0 (0.0%) | 0 (0.0%) | 1 (25.0%) | 17 (9.3%) | 4 (22.2%) | | |
| **Mild** | 18 (6.7%) | 0 (0.0%) | 0 (0.0%) | 0 (0.0%) | 0 (0.0%) | 2 (50.0%) | 15 (8.2%) | 1 (5.6%) | | |
| **Moderate** | 4 (1.5%) | 0 (0.0%) | 0 (0.0%) | 0 (0.0%) | 0 (0.0%) | 0 (0.0%) | 4 (2.2%) | 0 (0.0%) | | |
| **Severe** | 5 (1.9%) | 1 (2.3%) | 0 (0.0%) | 0 (0.0%) | 1 (20.0%) | 0 (0.0%) | 3 (1.6%) | 0 (0.0%) | | |
| **Subpleural scarring:** | | | | | | | | | 0.739 | 241 |
| **Nil** | 235 (97.5%) | 39 (97.5%) | 1 (100.0%) | 0 (.%) | 5 (100.0%) | 4 (100.0%) | 168 (97.1%) | 18 (100.0%) | | |
| **Trace** | 2 (0.8%) | 1 (2.5%) | 0 (0.0%) | 0 (.%) | 0 (0.0%) | 0 (0.0%) | 1 (0.6%) | 0 (0.0%) | | |
| **Mild** | 4 (1.7%) | 0 (0.0%) | 0 (0.0%) | 0 (.%) | 0 (0.0%) | 0 (0.0%) | 4 (2.3%) | 0 (0.0%) | | |

**Table V.** Number of PAH domain samples in the analysis broken down by maximal unrelated set and related set, and cases (included and excluded) and unaffected relatives.

| ***PAH domain samples*** | **Cases** | **Cases excluded** | **Unaffected relatives** | **Total** |
|---|---|---|---|---|
| **Max. unrelated set** | 1102 | 3 | 18 | **1123** |
| **Related set** | 20 | - | 5 | 25 |
| **Total** | **1122** | 3 | 23 | **1148** |

**Table VI.** Clinical characterization of the study population. Entire cohort (n=1,122) was composed of IPAH (n=972), HPAH (n=73), PVOD/PCH (n=20), PH associated with left heart disease (n=7), PH associated with lung disease (n=8), chronic thromboembolic pulmonary hypertension (n=6), multifactorial PH (n=6), hereditary hemorrhagic telangiectasia (n=1). Abbreviations: BMI - body mass index, WHO FC - World Health Organization Functional Class, 6MWD - 6-minute walk distance, mRAP - mean right atrial pressure, mPAP - mean pulmonary artery pressure, CO - cardiac output, FEV1 - forced expiratory capacity in 1 second, FVC - forced vital capacity, KCO - transfer factor coefficient for carbon monoxide, Hb - haemoglobin, RDW - red cell distribution width, WBC - white blood cell count, NTproBNP - N-terminal pro B-type natriuretic peptide, BNP - B-type natriuretic peptide, CRP - C-Reactive Protein Protein, HTN - hypertension, DM - diabetes mellitus, CAD - coronary artery disease, CVA - cerebrovascular accident, COPD - chronic obstructive pulmonary disease, CCB - calcium channel blocker, ERA - endothelin receptor antagonists, PA - prostacyclin analogues, PED5 - phosphodiesterase type 5, sGC - soluble guanylate cyclase.

| | ALL (N=1122) | N | I/HPAH and PVOD/PCH (N=1065) | N |
|---|---|---|---|---|
| **Demographics and functional status** | | | | |
| **Sex: female** | 760 (68%) | 1116 | 732 (69%) | 1064 |
| **Ethnicity: European** | 945 (84%) | 1122 | 898 (85%) | |
| **Age [years]** | 49 [35;63] | 1112 | 49 [35;63] | 1061 |
| **BMI [kg/m²]** | 27 [23;32] | 1015 | 27 [23;31] | 970 |
| **WHO FC: I/II/III/IV** | 21 (2%)/217 (20%)/703 (65%)/138 (13%) | 1079 | 21 (2%)/210 (20%)/663 (64%)/135 (13%) | 1029 |
| **6MWD [m]** | 335 [220;415] | 953 | 336 [220;415] | 906 |
| **Haemodynamics** | | | | |
| **mRAP [mmHg]** | 8 [5;12] | 985 | 8 [5;12] | 939 |
| **mPAP [mmHg]** | 53 [44;61] | 1052 | 53 [44;61] | 1004 |
| **CO [L/min]** | 3.9 [3.1;4.9] | 1003 | 3.9 [3.1;4.9] | 960 |
| **$FEV_1$ [% pred.]** | 85 [73;97] | 849 | 86 [74;97] | 811 |
| **FVC [% pred.]** | 94 [81;106] | 831 | 95 [82;106] | 793 |
| **KCO [% pred.]** | 71 [52;86] | 644 | 71 [52;86] | 610 |
| **Clinical blood tests** | | | | |
| **Hb [g/l]** | 151 [138;165] | 847 | 152 [138;164] | 805 |
| **RDW [%]** | 14 [14;16] | 413 | 14 [14;16] | 392 |
| **WBC [x10e9/l]** | 8.2 [6.8;9.8] | 839 | 8.2 [6.8;9.8] | 797 |
| **Platelets [x10e9/l]** | 224 [182;272] | 836 | 225 [183;274] | 795 |
| **Creatinine [µmol/l]** | 86 [70;102] | 832 | 86 [70;102] | 790 |
| **NTproBNP [ng/l]** | 926 [215;2637] | 276 | 963 [217;2672] | 265 |
| **BNP [ng/l]** | 195 [72;432] | 271 | 197 [74.6;454] | 252 |
| **CRP [mg/l]** | 4 [2;8] | 639 | 4 [2;8] | 604 |
| **Comorbidities** | | | | |

| | | | | |
|---|---|---|---|---|
| **HTN** | 265 (24%) | 1122 | 256 (24%) | 1065 |
| **DM type 1** | 20 (2%) | 1122 | 19 (2%) | 1065 |
| **DM type 2** | 138 (12%) | 1122 | 132 (12%) | 1065 |
| **CAD** | 45 (4%) | 1122 | 42 (4%) | 1065 |
| **CVA** | 17 (2%) | 1122 | 15 (1%) | 1065 |
| **Hypothyroidism** | 135 (12%) | 1122 | 130 (12%) | 1065 |
| **COPD** | 66 (6%) | 1122 | 57 (5%) | 1065 |
| **Asthma** | 78 (7%) | 1122 | 74 (7%) | 1065 |
| **Cancer** | 4 (0.4%) | 1122 | 3 (0.3%) | 1065 |
| **Medication** | | | | |
| **Initial therapy:** | | 631 | | 604 |
| **CCB** | 73 (12%) | | 72 (12%) | |
| **combination therapy** | 279 (44%) | | 269 (45%) | |
| **ERA** | 86 (14%) | | 82 (14%) | |
| **PA** | 42 (7%) | | 41 (7%) | |
| **PDE5 inhibitor** | 150 (24%) | | 139 (23%) | |
| **sGC stimulator** | 1 (0%) | | 1 (0%) | |

**Table VII.** Clinical differences between patients with present and absent transfer coefficient for carbon monoxide. Abbreviations: BMI - body mass index, WHO FC - World Health Organization Functional Class, 6MWD - 6-minute walk distance, mRAP - mean right atrial pressure, mPAP - mean pulmonary artery pressure, PAWP - Pulmonary Artery Wedge Pressure, CI - cardiac index, PVR - pulmonary vascular resistance, $SvO_2$ [%] - mixed venous saturation, HRCT - High-Resolution Computerized Tomography, FEV1 - forced expiratory capacity in 1 second, FVC - forced vital capacity, COPD - chronic obstructive pulmonary disease, OSA - obstructive sleep apnoea.

| | ALL (N=1122) | KCO [% pred.] missing (N=478) | KCO [% pred.] present (N=644) | p-value | N |
|---|---|---|---|---|---|
| **Sex: Female** | 760 (68.1%) | 326 (69.1%) | 434 (67.4%) | 0.597 | 1116 |
| **Prevalent cases** | 852 (75.9%) | 411 (86.0%) | 441 (68.5%) | <0.001 | 1122 |
| **BMI [kg/m²]** | 26.9 [23.1;31.5] | 25.6 [22.0;30.1] | 27.7 [24.1;32.5] | <0.001 | 1015 |
| **Age [years]** | 48.8 [34.8;62.7] | 44.6 [31.0;58.5] | 51.3 [38.1;65.5] | <0.001 | 1112 |
| **WHO FC** | | | | <0.001 | 1079 |
| **I** | 21 (1.95%) | 13 (2.88%) | 8 (1.27%) | | |
| **II** | 217 (20.1%) | 117 (25.9%) | 100 (15.9%) | | |
| **III** | 703 (65.2%) | 269 (59.6%) | 434 (69.1%) | | |
| **IV** | 138 (12.8%) | 52 (11.5%) | 86 (13.7%) | | |
| **6MWD [m]** | 340 [230;418] | 364 [289;432] | 314 [192;405] | <0.001 | 702 |
| **$FEV_1$ [% pred.]** | 85.0 [73.0;97.0] | 83.2 [69.6;97.0] | 86.0 [74.0;97.0] | 0.2 | 849 |
| **FVC [% pred.]** | 94.0 [81.4;106] | 89.0 [73.0;103] | 96.0 [82.6;107] | <0.001 | 831 |
| **TLC [% pred.]** | 95.0 [85.0;104] | 93.2 [83.7;104] | 95.0 [86.0;103] | 0.625 | 639 |
| **mRAP [mmHg]** | 8.00 [5.00;12.0] | 7.00 [5.00;12.0] | 9.00 [6.00;12.0] | <0.001 | 985 |
| **mPAP [mmHg]** | 53.0 [44.0;61.0] | 53.0 [44.0;61.0] | 53.0 [45.0;61.0] | 0.669 | 1052 |
| **PAWP [mmHg]** | 9.00 [7.00;12.0] | 9.00 [6.00;11.0] | 10.0 [7.00;12.0] | 0.001 | 934 |
| **CI [L/min/m²]** | 2.17 [1.72;2.67] | 2.31 [1.81;2.82] | 2.07 [1.68;2.59] | <0.001 | 946 |
| **PVR [WU]** | 11.0 [7.69;15.1] | 10.6 [7.60;15.2] | 11.1 [7.69;15.1] | 0.622 | 893 |
| **$SvO_2$ [%]** | 64.0 [58.0;70.0] | 64.7 [57.8;70.0] | 63.8 [58.2;70.0] | 0.732 | 817 |
| **Fibrosis [HRCT report]:** | | | | 0.445 | 614 |
| **none** | 586 (95.4%) | 169 (96.0%) | 417 (95.2%) | | |
| **minimal/mild** | 26 (4.23%) | 6 (3.41%) | 20 (4.57%) | | |
| **moderate** | 1 (0.16%) | 0 (0.00%) | 1 (0.23%) | | |
| **severe** | 1 (0.16%) | 1 (0.57%) | 0 (0.00%) | | |
| **Emphysema [HRCT report]:** | | | | 0.029 | 612 |
| **none** | 560 (91.5%) | 169 (96.0%) | 391 (89.7%) | | |

| | | | | | |
|---|---|---|---|---|---|
| **minimal/mild** | 33 (5.39%) | 3 (1.70%) | 30 (6.88%) | | |
| **moderate** | 15 (2.45%) | 4 (2.27%) | 11 (2.52%) | | |
| **severe** | 4 (0.65%) | 0 (0.00%) | 4 (0.92%) | | |
| **Smoking history: past/current** | 435 (50.6%) | 97 (38.5%) | 338 (55.7%) | <0.001 | 859 |
| **COPD** | 66 (5.88%) | 24 (5.02%) | 42 (6.52%) | 0.353 | 1122 |
| **OSA** | 58 (5.17%) | 20 (4.18%) | 38 (5.90%) | 0.251 | 1122 |
| **Asthma** | 78 (6.95%) | 15 (3.14%) | 63 (9.78%) | <0.001 | 1122 |

**Table VIII.** Clinical characteristics of unrelated individuals used in gene-tag association analysis by KCO threshold. None of the patients had systemic lupus erythematosus, systemic sclerosis, undifferentiated connective tissue disease or ankylosing spondylitis. Abbreviations: BMI - body mass index, WHO FC - World Health Organization Functional Class, 6MWD - 6-minute walk distance, SpO2 - peripheral capillary oxygen saturation, mRAP - mean right atrial pressure, mPAP - mean pulmonary artery pressure, PAWP - Pulmonary Artery Wedge Pressure, CO - cardiac output, SvO2 - Mixed venous oxygen saturation, FEV1 - forced expiratory capacity in 1 second, FVC - forced vital capacity, TLC - Total Lung Capacity, KCO - transfer coefficient for carbon monoxide, HRCT - High-Resolution Computerized Tomography, NTproBNP - N-terminal pro B-Type Natriuretic Peptide, BNP - B-Type Natriuretic Peptide, CRP - C-Reactive Protein Protein, Hb - hemoglobin, WBC - white blood cell count, COPD - chronic obstructive pulmonary disease, OSA - obstructive sleep apnoea, CAD - coronary artery disease, CVA - cerebrovascular accident, PAD - peripheral artery disease, HTN - hypertension, DM - diabetes mellitus.

| | **ALL (N=644)** | **KCO > 50% pred. (N=492)** | **KCO ≤ 50% pred. (N=152)** | **p-value** | **N** |
|---|---|---|---|---|---|
| **Age [years]** | 51 [38;66] | 47 [36;60] | 66 [54;71] | <0.001 | 644 |
| **Sex: female** | 434 (67%) | 352 (72%) | 82 (54%) | <0.001 | 644 |
| **Incident cases** | 203 (31.5%) | 134 (27.2%) | 69 (45.4%) | <0.001 | 644 |
| **BMI [kg/m²]** | 27.7 [24.1;32.5] | 27.7 [23.7;32.7] | 27.7 [24.8;32.3] | 0.806 | 629 |
| **WHO FC** | | | | 0.013 | 628 |
| **I** | 8 (1%) | 7 (1%) | 1 (1%) | | |
| **II** | 100 (16%) | 88 (18%) | 12 (8%) | | |
| **III** | 434 (69%) | 320 (67%) | 114 (77%) | | |
| **IV** | 86 (14%) | 64 (13%) | 22 (15%) | | |
| **6MWD [m]** | 313 [190;404] | 334 [229;414] | 219 [120;348] | <0.001 | 599 |
| **$SpO_2$ pre [%]** | 95.0 [93.0;97.0] | 96.0 [93.0;98.0] | 92.0 [89.0;95.0] | <0.001 | 575 |
| **$SpO_2$ post [%]** | 91.0 [85.0;96.0] | 93.0 [88.0;96.0] | 83.0 [76.0;88.0] | <0.001 | 529 |
| **mRAP [mmHg]** | 9 [6;12] | 9 [6;13] | 9 [6;12] | 0.375 | 601 |
| **mPAP [mmHg]** | 53 [45;61] | 54 [46;63] | 50 [42;57] | 0.001 | 625 |
| **PAWP [mmHg]** | 10 [7;12] | 10 [7;12] | 10 [8;12] | 0.301 | 560 |
| **CO [L/min]** | 3.8 [3.1;4.8] | 3.9 [3.1;4.9] | 3.6 [3.1;4.7] | 0.282 | 614 |
| **$SvO_2$ [%]** | 64 [58;70] | 64 [59;71] | 61 [55;67] | <0.001 | 572 |
| **Acute NO challenge: vasoresponder** | 43 (17%) | 38 (18%) | 5 (10%) | 0.204 | 257 |
| **$FEV_1$ [% pred.]** | 86.0 [74.0;97.0] | 85.0 [73.4;96.0] | 87.0 [77.0;98.0] | 0.119 | 639 |
| **FVC [% pred.]** | 96.0 [82.6;107] | 94.0 [81.0;106] | 101 [87.0;113] | <0.001 | 628 |
| **$FEV_1$/FVC ratio** | 0.76 [0.69;0.81] | 0.76 [0.71;0.81] | 0.70 [0.63;0.77] | <0.001 | 614 |
| **TLC [% pred.]** | 95.0 [86.0;103] | 95.0 [85.0;103] | 95.0 [87.5;104] | 0.564 | 485 |
| **KCO [%pred.]** | 71 [52;86] | 78 [67;90] | 37 [30;44] | <0.001 | 644 |
| **Emphysema (HRCT scan)** | | | | <0.001 | 436 |

| | | | | | |
|---|---|---|---|---|---|
| **none** | 391 (90%) | 307 (95%) | 84 (74%) | | |
| **minimal/mild** | 30 (7%) | 12 (4%) | 18 (16%) | | |
| **moderate** | 11 (3%) | 2 (1%) | 9 (8%) | | |
| **severe** | 4 (1%) | 1 (0%) | 3 (3%) | | |
| **Fibrosis (HRCT scan)** | | | | <0.001 | 438 |
| **none** | 417 (95%) | 319 (98%) | 98 (87%) | | |
| **minimal/mild** | 20 (5%) | 6 (2%) | 14 (12%) | | |
| **moderate** | 1 (0%) | 0 (0%) | 1 (1%) | | |
| **Smoking history: past/current** | 338 (55.7%) | 233 (50.0%) | 105 (74.5%) | <0.001 | 607 |
| **NTproBNP [ng/l]** | 980 [248;2673] | 866 [257;2590] | 1225 [249;2878] | 0.359 | 220 |
| **BNP [ng/l]** | 200 [72.9;432] | 198 [69.7;456] | 200 [82.2;328] | 0.798 | 143 |
| **Uric acid [mmol/l]** | 0.42 [0.32;0.53] | 0.40 [0.30;0.51] | 0.48 [0.39;0.56] | 0.002 | 231 |
| **CRP [mg/l]** | 5.00 [2.00;8.60] | 5.00 [2.00;8.00] | 4.15 [2.00;9.00] | 0.953 | 450 |
| **Hb [g/l]** | 154 [139;166] | 153 [138;166] | 154 [142;166] | 0.658 | 603 |
| **WBC [x10e9/l]** | 8 [7;10] | 8 [7;10] | 9 [7;10] | 0.011 | 597 |
| **Platelets [x10e9/l]** | 220 [179;268] | 220 [179;270] | 220 [182;254] | 0.516 | 595 |
| **Sodium [mmol/l]** | 139 [138;141] | 139 [138;141] | 140 [138;141] | 0.858 | 597 |
| **Potassium [mmol/l]** | 4.20 [4.00;4.50] | 4.20 [3.90;4.50] | 4.20 [4.00;4.50] | 0.412 | 591 |
| **Urea [mmol/l]** | 5.80 [4.60;7.70] | 5.60 [4.40;7.10] | 7.00 [5.30;9.20] | <0.001 | 594 |
| **Creatinine [µmol/l]** | 88.0 [73.2;105] | 87.0 [73.0;102] | 95.0 [78.5;114] | 0.002 | 598 |
| **COPD** | 42 (7%) | 23 (5%) | 19 (12%) | 0.001 | 644 |
| **Asthma** | 63 (10%) | 54 (11%) | 9 (6%) | 0.093 | 644 |
| **OSA** | 38 (6%) | 29 (6%) | 9 (6%) | 1 | 644 |
| **CAD** | 26 (4%) | 10 (2%) | 16 (11%) | <0.001 | 644 |
| **CVA** | 10 (2%) | 5 (1%) | 5 (3%) | 0.061 | 644 |
| **PAD** | 2 (0%) | 0 (0%) | 2 (1%) | 0.055 | 644 |
| **HTN** | 170 (26%) | 116 (24%) | 54 (36%) | 0.005 | 644 |
| **DM type 1** | 9 (1%) | 8 (2%) | 1 (1%) | 0.693 | 644 |
| **DM type 2** | 93 (14%) | 60 (12%) | 33 (22%) | 0.005 | 644 |
| **Hypothyroidism** | 73 (11%) | 60 (12%) | 13 (9%) | 0.275 | 644 |
| **Sjogren syndrome** | 3 (0%) | 2 (0%) | 1 (1%) | 0.555 | 644 |

**Table IX.** Clinical characteristics of study individuals by KCO tertiles. None of the patients had systemic lupus erythematosus, systemic sclerosis, undifferentiated connective tissue disease or ankylosing spondylitis Abbreviations: BMI - body mass index, WHO FC - World Health Organisation Functional Class, 6MWD - 6-minute walk distance, $SpO_2$ - peripheral capillary oxygen saturation, mRAP - mean right atrial pressure, mPAP - mean pulmonary artery pressure, PAWP - Pulmonary Artery Wedge Pressure, CO - cardiac output, $SvO_2$ - mixed venous oxygen saturation, NO - nitric oxide, FEV1 - forced expiratory capacity in 1 second, FVC - forced vital capacity, TLC - Total Lung Capacity, KCO - transfer coefficient for carbon monoxide, HRCT - High-Resolution Computerized Tomography, NTproBNP - N-terminal pro B-type natriuretic peptide, BNP - B-type natriuretic peptide, CRP - C-Reactive Protein Protein, Hb - haemoglobin, WBC - white blood cell count, COPD - chronic obstructive pulmonary disease, OSA - obstructive sleep apnoea, CAD - coronary artery disease, CVA - cerebrovascular accident, PAD - peripheral artery disease, HTN - hypertension, DM - diabetes mellitus.

| | ALL (N=644) | Higher tertile (N=214) | Middle tertile (N=215) | Lower tertile (N=215) | p.overall | N |
|---|---|---|---|---|---|---|
| **Age [years]** | 51 [38;66] | 44 [37;58] | 49 [34;61] | 64 [50;71] | <0.001 | 644 |
| **Sex: female** | 434 (67%) | 149 (70%) | 160 (74%) | 125 (58%) | 0.001 | 644 |
| **Incident cases** | 203 (31.5%) | 58 (27.1%) | 54 (25.1%) | 91 (42.3%) | <0.001 | 644 |
| **BMI [kg/m²]** | 27.7 [24.1;32.5] | 29.1 [25.3;34.5] | 26.8 [23.1;30.9] | 27.4 [24.2;32.3] | <0.001 | 629 |
| **WHO FC** | | | | | 0.028 | 628 |
| **I** | 8 (1%) | 4 (2%) | 2 (1%) | 2 (1%) | | |
| **II** | 100 (16%) | 40 (19%) | 40 (19%) | 20 (10%) | | |
| **III** | 434 (69%) | 143 (68%) | 135 (65%) | 156 (74%) | | |
| **IV** | 86 (14%) | 22 (11%) | 32 (15%) | 32 (15%) | | |
| **6MWD [m]** | 313 [190;404] | 331 [240;420] | 340 [230;418] | 240 [131;360] | <0.001 | 599 |
| **$SpO_2$ pre [%]** | 95.0 [93.0;97.0] | 96.0 [94.0;98.0] | 96.0 [93.0;98.0] | 93.0 [90.0;96.0] | <0.001 | 575 |
| **$SpO_2$ post [%]** | 91.0 [85.0;96.0] | 94.0 [89.0;96.0] | 93.0 [88.0;96.0] | 85.0 [79.0;91.0] | <0.001 | 529 |
| **mRAP [mmHg]** | 9 [6;12] | 9 [7;13] | 9 [6;12] | 8 [5;12] | 0.224 | 601 |
| **mPAP [mmHg]** | 53 [45;61] | 55 [47;65] | 53 [46;62] | 51 [42;57] | <0.001 | 625 |
| **PAWP [mmHg]** | 10 [7;12] | 10 [8;12] | 10 [7;12] | 10 [7;12] | 0.487 | 560 |
| **CO [L/min]** | 3.8 [3.1;4.8] | 3.8 [3.0;5.0] | 4.0 [3.1;4.8] | 3.8 [3.1;4.8] | 0.967 | 614 |
| **$SvO_2$ [%]** | 64 [58;70] | 64 [60;71] | 65 [59;70] | 62 [56;68] | 0.001 | 572 |
| **Acute NO challenge: vasoresponder** | 43 (17%) | 15 (17%) | 21 (24%) | 7 (9%) | 0.024 | 257 |
| **$FEV_1$ [% pred.]** | 86.0 [74.0;97.0] | 83.8 [72.9;95.0] | 86.0 [73.9;97.4] | 87.0 [77.0;98.0] | 0.08 | 639 |
| **FVC [% pred.]** | 96.0 [82.6;107] | 90.0 [79.8;101] | 95.9 [83.0;108] | 100 [86.9;112] | <0.001 | 628 |
| **$FEV_1$/FVC ratio** | 0.76 [0.69;0.81] | 0.78 [0.72;0.82] | 0.76 [0.69;0.81] | 0.71 [0.65;0.78] | <0.001 | 614 |
| **TLC [% pred.]** | 95.0 [86.0;103] | 94.0 [86.2;103] | 96.0 [83.0;104] | 97.0 [87.0;102] | 0.787 | 485 |
| **KCO [% pred.]** | 71 [52;86] | 92 [86;101] | 71 [67;76] | 42 [33;52] | <0.001 | 644 |
| **Emphysema (HRCT scan):** | | | | | <0.001 | 436 |
| **none** | 391 (90%) | 144 (99%) | 128 (96%) | 119 (76%) | | |
| **minimal/mild** | 30 (7%) | 2 (1%) | 4 (3%) | 24 (15%) | | |

| | | | | | | |
|---|---|---|---|---|---|---|
| **moderate** | 11 (3%) | 0 (0%) | 0 (0%) | 11 (7%) | | |
| **severe** | 4 (1%) | 0 (0%) | 1 (1%) | 3 (2%) | | |
| **Fibrosis (HRCT scan):** | | | | | <0.001 | 438 |
| **none** | 417 (95%) | 145 (99%) | 132 (99%) | 140 (89%) | | |
| **minimal/mild** | 20 (5%) | 2 (1%) | 2 (1%) | 16 (10%) | | |
| **moderate** | 1 (0%) | 0 (0%) | 0 (0%) | 1 (1%) | | |
| **Smoking history: past/current** | 338 (55.7%) | 89 (42.4%) | 106 (52.7%) | 143 (73.0%) | <0.001 | 607 |
| **NTproBNP [ng/l]** | 980 [248;2673] | 842 [167;2358] | 902 [310;2593] | 1225 [237;2811] | 0.539 | 220 |
| **BNP [ng/l]** | 200 [72.9;432] | 193 [71.7;392] | 145 [67.5;427] | 214 [82.2;448] | 0.659 | 143 |
| **Uric acid [mmol/l]** | 0.42 [0.32;0.53] | 0.40 [0.28;0.47] | 0.40 [0.32;0.51] | 0.48 [0.38;0.55] | 0.011 | 231 |
| **CRP [mg/l]** | 5.00 [2.00;8.60] | 5.00 [2.00;8.00] | 5.00 [2.00;9.32] | 4.00 [2.00;8.57] | 0.496 | 450 |
| **Hb [g/l]** | 154 [139;166] | 160 [145;169] | 149 [136;164] | 151 [140;165] | <0.001 | 603 |
| **WBC [x10e9/l]** | 8 [7;10] | 8 [7;9] | 8 [7;10] | 9 [7;10] | 0.005 | 597 |
| **Platelets [x10e9/l]** | 220 [179;268] | 224 [183;262] | 219 [174;276] | 217 [184;262] | 0.89 | 595 |
| **Sodium [mmol/l]** | 139 [138;141] | 140 [138;141] | 139 [137;141] | 140 [138;141] | 0.623 | 597 |
| **Potassium [mmol/l]** | 4.20 [4.00;4.50] | 4.30 [4.00;4.50] | 4.20 [3.90;4.40] | 4.30 [4.00;4.50] | 0.13 | 591 |
| **Urea [mmol/l]** | 5.80 [4.60;7.70] | 5.50 [4.43;7.00] | 5.70 [4.30;7.10] | 6.70 [5.15;8.80] | <0.001 | 594 |
| **Creatinine [µmol/l]** | 88.0 [73.2;105] | 87.0 [73.0;99.5] | 86.0 [72.0;102] | 92.5 [77.0;111] | 0.003 | 598 |
| **COPD** | 42 (7%) | 5 (2%) | 12 (6%) | 25 (12%) | <0.001 | 644 |
| **Asthma** | 63 (10%) | 26 (12%) | 25 (12%) | 12 (6%) | 0.039 | 644 |
| **OSA** | 38 (6%) | 12 (6%) | 11 (5%) | 15 (7%) | 0.698 | 644 |
| **CAD** | 26 (4%) | 2 (1%) | 4 (2%) | 20 (9%) | <0.001 | 644 |
| **CVA** | 10 (2%) | 1 (0%) | 3 (1%) | 6 (3%) | 0.174 | 644 |
| **PAD** | 2 (0%) | 0 (0%) | 0 (0%) | 2 (1%) | 0.332 | 644 |
| **HTN** | 170 (26%) | 46 (21%) | 53 (25%) | 71 (33%) | 0.02 | 644 |
| **DM type 1** | 9 (1%) | 2 (1%) | 5 (2%) | 2 (1%) | 0.526 | 644 |
| **DM type 2** | 93 (14%) | 22 (10%) | 22 (10%) | 49 (23%) | <0.001 | 644 |
| **Hypothyroidism** | 73 (11%) | 25 (12%) | 30 (14%) | 18 (8%) | 0.185 | 644 |
| **Sjogren syndrome** | 3 (0%) | 2 (1%) | 0 (0%) | 1 (0%) | 0.331 | 644 |

**Table X.** Results of Cox regression analysis relating overall survival to selected variables at baseline. Abbreviations: CI - Confidence interval, 6MWD - 6-minute walking distance, mPAP - mean pulmonary arterial pressure, mRAP - mean right atrial pressure, PVR - pulmonary vascular resistance, WU - Wood units, KCO - transfer coefficient for carbon monoxide, CAD - coronary artery disease, COPD - chronic obstructive pulmonary disease, HTN - systemic hypertension, HRCT - High-Resolution Computerized Tomography.

| | | | Univariate | | Multivariate | |
|---|---|---|---|---|---|---|
| | No event (N=481) | Event (N=163) | HR [95% CI] | p-value | HR [95% CI] | p-value |
| **Sex:** | | | | <0.001 | | <0.001 |
| **female** | 343 (71.3%) | 91 (55.8%) | Ref. | | Ref. | |
| **male** | 138 (28.7%) | 72 (44.2%) | 1.98 [1.45;2.70] | | 2.93 [1.81;4.75] | |
| **Age [years]** | 4.80 (1.50) | 6.17 (1.59) | 1.75 [1.56;1.96] | <0.001 | 1.57 [1.27;1.94] | <0.001 |
| **Incident/Prevalent:** | | | | <0.001 | | 0.131 |
| **incident** | 151 (31.4%) | 52 (31.9%) | Ref. | | Ref. | |
| **prevalent** | 330 (68.6%) | 111 (68.1%) | 0.40 [0.28;0.58] | | 0.65 [0.37;1.14] | |
| **6MWD [m]** | 32.9 (15.0) | 21.7 (12.9) | 0.95 [0.94;0.96] | <0.001 | 0.97 [0.95;0.99] | 0.002 |
| **mRAP [mmHg]** | 1.84 (1.07) | 2.15 (1.13) | 1.26 [1.11;1.44] | 0.001 | 1.28 [1.05;1.57] | 0.016 |
| **mPAP [mmHg]** | 10.9 (2.75) | 10.4 (2.34) | 0.94 [0.88;1.00] | 0.038 | 1.08 [0.97;1.2] | 0.142 |
| **CI [L/min/m²]** | 2.28 (0.78) | 2.10 (0.68) | 0.67 [0.53;0.86] | 0.002 | 0.98 [0.62;1.55] | 0.923 |
| **PVR [WU]** | 12.1 (5.96) | 11.8 (4.94) | 1.00 [0.97;1.03] | 0.89 | | |
| **KCO [%pred]** | 7.35 (2.22) | 5.71 (2.41) | 0.71 [0.66;0.77] | <0.001 | 0.79 [0.7;0.88] | <0.001 |
| **Smoking history:** | | | | 0.002 | | 0.692 |
| **no** | 215 (47.5%) | 54 (35.1%) | Ref. | | Ref. | |
| **past/current** | 238 (52.5%) | 100 (64.9%) | 1.67 [1.20;2.33] | | 1.11 [0.66;1.88] | |
| **CAD:** | | | | 0.002 | | 0.081 |
| **no** | 467 (97.1%) | 151 (92.6%) | Ref. | | Ref. | |
| **yes** | 14 (2.91%) | 12 (7.36%) | 2.51 [1.39;4.53] | | 0.38 [0.13;1.13] | |
| **COPD:** | | | | 0.141 | | 0.268 |
| **no** | 452 (94.0%) | 150 (92.0%) | Ref. | | Ref. | |
| **yes** | 29 (6.03%) | 13 (7.98%) | 1.53 [0.87;2.69] | | 0.59 [0.23;1.5] | |
| **HTN:** | | | | <0.001 | | |
| **no** | 374 (77.8%) | 100 (61.3%) | Ref. | | Ref. | |
| **yes** | 107 (22.2%) | 63 (38.7%) | 1.92 [1.40;2.63] | | 1.12 [0.68;1.84] | |
| **Emphysema (HRCT scan):** | | | | 0.012 | | |
| **none** | 300 (91.5%) | 91 (84.3%) | Ref. | | Ref. | |
| **minimal/mild** | 19 (5.79%) | 11 (10.2%) | 2.12 [1.13;3.98] | | 0.85 [0.38;1.92] | 0.696 |

| | | | | | | |
|---|---|---|---|---|---|---|
| **moderate** | 6 (1.83%) | 5 (4.63%) | 2.83 [1.15;6.98] | | 0.7 [0.19;2.61] | 0.594 |
| **severe** | 3 (0.91%) | 1 (0.93%) | 2.36 [0.33;17.0] | | 0 [0;Inf] | 0.996 |
| **Fibrosis (HRCT scan):** | | | | 0.003 | | |
| **none** | 321 (97.0%) | 96 (89.7%) | Ref. | | Ref. | |
| **minimal/mild** | 10 (3.02%) | 10 (9.35%) | 2.79 [1.45;5.36] | | 0.83 [0.36;1.93] | 0.672 |
| **moderate** | 0 (0.00%) | 1 (0.93%) | 3.29 [0.46;23.6] | | 3.98 [0.47;33.58] | 0.204 |

**Table XI.** Clinical characteristics of all unrelated individuals used in gene-tag association by age tertiles. None of the patients had undifferentiated connective tissue disease, incident cases were defined as those diagnosed within 6 months from study commencement. Abbreviations: BMI - body mass index, WHO FC - World Health Organization Functional Class, 6MWD - 6-minute walk distance, $SpO_2$ - peripheral capillary oxygen saturation, mRAP - mean right atrial pressure, mPAP - mean pulmonary artery pressure, PAWP - Pulmonary Artery Wedge Pressure, CO - cardiac output, $SvO_2$ - Mixed venous oxygen saturation, NO - nitric oxide, FEV1 - forced expiratory capacity in 1 second, FVC - forced vital capacity, TLC - Total Lung Capacity, KCO - transfer factor coefficient for carbon monoxide, NTproBNP - N-terminal pro B-type natriuretic peptide, BNP - B-type natriuretic peptide, CRP - C-Reactive Protein Protein, Hb - hemoglobin, WBC - white blood cell count, COPD - chronic obstructive pulmonary disease, OSA - obstructive sleep apnoea, CAD - coronary artery disease, CVA - cerebrovascular accident, PAD - peripheral artery disease, HTN - hypertension, DM - diabetes mellitus.

| | ALL (N=1112) | Lower tertile (N=381) | Middle tertile (N=376) | Higher tertile (N=355) | p.overall | N |
|---|---|---|---|---|---|---|
| **Age [years]** | 49 [35;63] | 30 [25;35] | 50 [45;54] | 68 [64;73] | <0.001 | 1112 |
| **Sex: female** | 757 (68%) | 276 (72%) | 273 (73%) | 208 (59%) | <0.001 | 1112 |
| **Incident cases** | 270 (24.3%) | 77 (20.2%) | 74 (19.7%) | 119 (33.5%) | <0.001 | 1112 |
| **BMI [kg/m²]** | 26.9 [23.1;31.5] | 24.2 [21.0;29.3] | 28.0 [24.3;32.5] | 28.1 [25.1;32.0] | <0.001 | 1015 |
| **WHO FC** | | | | | <0.001 | 1078 |
| **I** | 21 (2%) | 15 (4%) | 3 (1%) | 3 (1%) | | |
| **II** | 217 (20%) | 95 (26%) | 79 (21%) | 43 (12%) | | |
| **III** | 703 (65%) | 201 (56%) | 248 (67%) | 254 (73%) | | |
| **IV** | 137 (13%) | 50 (14%) | 40 (11%) | 47 (14%) | | |
| **6MWD [m]** | 335 [220;415] | 375 [302;460] | 343 [250;420] | 236 [134;348] | <0.001 | 953 |
| **$SpO_2$ pre [%]** | 96.0 [93.0;97.0] | 97.0 [95.0;98.0] | 96.0 [93.0;97.0] | 94.0 [90.0;96.0] | <0.001 | 890 |
| **$SpO_2$ post [%]** | 91.0 [85.0;95.0] | 94.0 [88.0;97.0] | 92.0 [86.0;96.0] | 88.0 [82.0;92.8] | <0.001 | 830 |
| **mRAP [mmHg]** | 8 [5;12] | 8 [5;12] | 9 [6;13] | 8 [5;12] | 0.046 | 984 |
| **mPAP [mmHg]** | 53 [44;61] | 55 [47;66] | 55 [48;62] | 48 [40;57] | <0.001 | 1051 |
| **PAWP [mmHg]** | 9 [7;12] | 9 [6;11] | 9 [7;12] | 10 [7;13] | 0.004 | 933 |
| **CO [L/min]** | 3.9 [3.1;4.9] | 4.0 [3.1;5.0] | 3.9 [3.1;4.9] | 3.8 [3.2;4.8] | 0.651 | 1003 |
| **$SvO_2$ [%]** | 64 [58;70] | 67 [60;72] | 64 [58;70] | 62 [57;67] | <0.001 | 817 |
| **Acute NO challenge** | 59 (14%) | 31 (17%) | 21 (14%) | 7 (7%) | 0.078 | 435 |
| **$FEV_1$ [% pred.]** | 85.0 [73.0;97.0] | 87.0 [77.0;97.0] | 84.0 [71.0;96.0] | 85.2 [71.1;97.9] | 0.34 | 849 |
| **FVC [% pred.]** | 94.0 [81.4;106] | 90.0 [81.0;102] | 96.0 [82.0;108] | 96.3 [82.0;108] | 0.018 | 831 |
| **$FEV_1$/FVC ratio** | 0.76 [0.69;0.81] | 0.81 [0.77;0.86] | 0.75 [0.69;0.80] | 0.72 [0.66;0.77] | <0.001 | 760 |
| **TLC [% pred.]** | 95.0 [85.0;104] | 95.0 [85.0;103] | 95.0 [87.0;106] | 93.6 [83.0;102] | 0.082 | 639 |
| **KCO [% pred.]** | 71 [52;86] | 76 [65;91] | 76 [62;88] | 57 [40;78] | <0.001 | 644 |
| **Emphysema:** | | | | | <0.001 | 611 |
| **none** | 559 (91%) | 193 (99%) | 198 (93%) | 168 (82%) | | |
| **minimal/mild** | 33 (5%) | 1 (1%) | 10 (5%) | 22 (11%) | | |

| | | | | | | |
|---|---|---|---|---|---|---|
| **moderate** | 15 (2%) | 0 (0%) | 4 (2%) | 11 (5%) | | |
| **severe** | 4 (1%) | 0 (0%) | 0 (0%) | 4 (2%) | | |
| **Fibrosis:** | | | | | <0.001 | 613 |
| **none** | 585 (95%) | 193 (98%) | 208 (98%) | 184 (90%) | | |
| **minimal/mild** | 26 (4%) | 3 (2%) | 2 (1%) | 21 (10%) | | |
| **moderate** | 1 (0%) | 0 (0%) | 1 (0%) | 0 (0%) | | |
| **severe** | 1 (0%) | 0 (0%) | 1 (0%) | 0 (0%) | | |
| **Smoking history: past/current** | 435 (50.8%) | 107 (36.9%) | 153 (53.5%) | 175 (62.3%) | <0.001 | 857 |
| **NTproBNP [ng/l]** | 926 [215;2637] | 345 [122;1640] | 763 [158;1356] | 1996 [501;3706] | <0.001 | 276 |
| **BNP [ng/l]** | 195 [72.4;432] | 117 [30.0;394] | 181 [85.1;398] | 236 [112;481] | 0.005 | 271 |
| **Uric acid [mmol/l]** | 0.41 [0.31;0.52] | 0.37 [0.26;0.46] | 0.41 [0.30;0.50] | 0.48 [0.36;0.56] | <0.001 | 358 |
| **CRP [mg/l]** | 4.30 [2.00;8.50] | 4.00 [2.00;7.00] | 4.15 [2.00;8.50] | 5.00 [2.50;9.10] | 0.151 | 639 |
| **Hb [g/l]** | 151 [138;165] | 152 [138;164] | 154 [141;166] | 149 [133;163] | 0.007 | 847 |
| **WBC [x10e9/l]** | 8 [7;10] | 8 [6;10] | 8 [7;10] | 8 [7;10] | 0.73 | 839 |
| **Platelets [x10e9/l]** | 224 [182;272] | 231 [186;280] | 219 [181;261] | 221 [179;272] | 0.072 | 836 |
| **Sodium [mmol/l]** | 140 [138;141] | 140 [138;141] | 139 [138;141] | 140 [137;141] | 0.728 | 835 |
| **Potassium [mmol/l]** | 4.20 [3.90;4.50] | 4.20 [3.98;4.40] | 4.20 [3.90;4.40] | 4.30 [4.00;4.50] | 0.03 | 830 |
| **Urea [mmol/l]** | 5.70 [4.40;7.60] | 4.80 [3.88;5.81] | 5.50 [4.30;6.70] | 7.60 [5.90;10.1] | <0.001 | 830 |
| **Creatinine [µmol/l]** | 85.5 [70.0;102] | 79.0 [68.0;93.5] | 82.0 [69.0;96.0] | 96.0 [80.0;121] | <0.001 | 832 |
| **COPD** | 65 (6%) | 2 (1%) | 22 (6%) | 41 (12%) | <0.001 | 1112 |
| **Asthma** | 78 (7%) | 32 (8%) | 32 (9%) | 14 (4%) | 0.023 | 1112 |
| **OSA** | 57 (5%) | 5 (1%) | 21 (6%) | 31 (9%) | <0.001 | 1112 |
| **CAD** | 44 (4%) | 0 (0%) | 9 (2%) | 35 (10%) | <0.001 | 1112 |
| **CVA** | 17 (2%) | 2 (1%) | 6 (2%) | 9 (3%) | 0.084 | 1112 |
| **PAD** | 5 (0%) | 0 (0%) | 0 (0%) | 5 (1%) | 0.003 | 1112 |
| **HTN** | 264 (24%) | 19 (5%) | 80 (21%) | 165 (46%) | <0.001 | 1112 |
| **DM type 1** | 19 (2%) | 7 (2%) | 6 (2%) | 6 (2%) | 0.967 | 1112 |
| **DM type 2** | 137 (12%) | 5 (1%) | 37 (10%) | 95 (27%) | <0.001 | 1112 |
| **Hypothyroidism** | 135 (12%) | 38 (10%) | 49 (13%) | 48 (14%) | 0.274 | 1112 |
| **Systemic lupus erythormatosus** | 1 (0%) | 0 (0%) | 0 (0%) | 1 (0%) | 0.319 | 1112 |
| **Systemic sclerosis** | 1 (0%) | 0 (0%) | 0 (0%) | 1 (0%) | 0.319 | 1112 |
| **Ankylosing spondylitis** | 1 (0.09%) | 0 (0.00%) | 0 (0.00%) | 1 (0.28%) | 0.319 | 1112 |
| **Sjogren syndrome** | 5 (0%) | 2 (1%) | 1 (0%) | 2 (1%) | 0.871 | 1112 |

**Table XII.** Clinical characteristics of IPAH patients who harbor loss-of-function variants in *BMPR2*, *EIF2AK4* and *KDR*. Comorbidities are reported as the number and percentage of cases possessing a disease entity. Abbreviations: BMI - body mass index, WHO FC - World Health Organization functional class, 6MWD - 6-minute walk distance, SpO2 - arterial oxygen saturation, mRAP - mean right atrial pressure, mPAP - mean pulmonary artery pressure, mPAWP - mean pulmonary artery wedge pressure, CO - cardiac output, PVR - pulmonary vascular resistance, NO - nitric oxide challenge, FEV1 - forced expiratory volume in 1 second, FVC - forced vital capacity, KCO - transfer factor coefficient for carbon monoxide, COPD - chronic obstructive pulmonary disease, OSA - obstructive sleep apnea, CAD - coronary artery disease, HTN - systemic hypertension, CKD - chronic kidney disease, Hb - hemoglobin, WBC - white blood cells, TSH - thyroid-stimulating hormone.

| | ***BMPR2*** **(N=162)** | ***EIF2AK4*** **(bial.) (N=14)** | ***KDR*** **(lof) (N=4)** | **No mutation (N=829)** | **p.overall** | **N** |
|---|---|---|---|---|---|---|
| **Age[years]** | 39 [32;51] | 31 [23;42] | 64 [62;68] | 51 [38;66] | <0.001 | 1000 |
| **Sex: female** | 107 (66%) | 7 (50%) | 2 (50%) | 579 (70%) | 0.184 | 1004 |
| **BMI[kg/m^2]** | 27 [23;32] | 24 [20;27] | 26 [26;30] | 27 [23;32] | 0.07 | 915 |
| **WHO functional class:** | | | | | 0.334 | 970 |
| **I** | 2 (1.2%) | 0 (0.0%) | 0 (0.0%) | 17 (2.1%) | | |
| **II** | 32 (19.9%) | 2 (14.3%) | 1 (25.0%) | 154 (19.5%) | | |
| **III** | 96 (59.6%) | 9 (64.3%) | 3 (75.0%) | 528 (66.8%) | | |
| **IV** | 31 (19.3%) | 3 (21.4%) | 0 (0.0%) | 92 (11.6%) | | |
| **6MWD[m]** | 355 [288;421] | 302 [210;466] | 301 [240;362] | 332 [218;412] | 0.27 | 628 |
| **SpO2 pre** | 96 [94;98] | 92 [90;96] | 97 [96;97] | 95 [93;97] | 0.003 | 804 |
| **SpO2 post** | 94 [89;97] | 83 [76;86] | 86 [86;88] | 91 [84;95] | <0.001 | 746 |
| **mRAP[mmHg]** | 10 [6;14] | 8 [6;10] | 6 [4;8] | 8 [5;12] | 0.013 | 886 |
| **mPAP[mmHg]** | 57 [52;67] | 52 [44;59] | 50 [43;58] | 52 [42;61] | <0.001 | 950 |
| **mPAWP[mmHg]** | 10 [7;12] | 11 [8;12] | 10 [8;13] | 9 [7;12] | 0.864 | 842 |
| **CO[L/min]** | 3.3 [2.7;3.9] | 4.5 [3.0;4.9] | 4.9 [4.3;5.4] | 4.0 [3.2;5.1] | <0.001 | 908 |
| **PVR[WU]** | 14.5 [10.8;20.4] | 9.56 [8.16;11.1] | 7.93 [6.23;10.9] | 10.3 [7.13;13.9] | <0.001 | 808 |
| **Acute NO challenge: vasoresponder** | 1 (1.28%) | 0 (0.00%) | 1 (25.0%) | 53 (17.5%) | <0.001 | 392 |
| **FEV1[% pred.]** | 91 [79;100] | 93 [84;100] | 86 [79;96] | 84 [71;95] | <0.001 | 767 |
| **FVC[% pred.]** | 100 (17) | 101 (16) | 93 (17) | 92 (20) | 0.001 | 751 |
| **FEV1/FVC ratio** | 0.77 [0.73;0.82] | 0.79 [0.69;0.81] | 0.78 [0.76;0.79] | 0.76 [0.69;0.81] | 0.063 | 685 |
| **KCO [% pred]** | 83 [74;96] | 33 [30;35] | 46 [46;48] | 68 [48;83] | <0.001 | 580 |
| **Smoking: current/previous** | 53 (40.8%) | 4 (30.8%) | 1 (25.0%) | 334 (53.2%) | 0.015 | 775 |
| **COPD: yes** | 6 (3.70%) | 0 (0.00%) | 0 (0.00%) | 49 (5.91%) | 0.6 | 1009 |
| **Asthma: yes** | 20 (12.3%) | 4 (28.6%) | 0 (0.00%) | 47 (5.67%) | 0.001 | 1009 |
| **OSA: yes** | 0 (0.00%) | 0 (0.00%) | 0 (0.00%) | 56 (6.76%) | 0.001 | 1009 |
| **CAD: yes** | 3 (1.85%) | 1 (7.14%) | 1 (25.0%) | 32 (3.86%) | 0.071 | 1009 |
| **HTN: yes** | 28 (17.3%) | 0 (0.00%) | 2 (50.0%) | 213 (25.7%) | 0.005 | 1009 |

| | | | | | | |
|---|---|---|---|---|---|---|
| **CKD: yes** | 4 (2.47%) | 0 (0.00%) | 0 (0.00%) | 43 (5.19%) | 0.418 | 1009 |
| **Hypothyroidism: yes** | 14 (8.64%) | 1 (7.14%) | 1 (25.0%) | 108 (13.0%) | 0.251 | 1009 |
| **Pulmonary embolism: yes** | 2 (1.23%) | 0 (0.00%) | 0 (0.00%) | 27 (3.26%) | 0.482 | 1009 |
| **Hb[g/l]** | 162 [152;173] | 165 [154;179] | 148 [135;152] | 149 [135;162] | <0.001 | 764 |
| **WBC[x10e9/l]** | 8.74 [7.30;10.8] | 7.43 [6.50;10.8] | 8.80 [8.23;9.55] | 8.10 [6.70;9.70] | 0.019 | 757 |
| **Platelets[x10e9/l]** | 210 [174;251] | 219 [206;234] | 216 [188;251] | 228 [181;277] | 0.268 | 753 |
| **TSH[mU/l]** | 2.37 [1.67;3.65] | 2.08 [1.09;3.69] | 1.76 [1.72;1.84] | 2.00 [1.10;3.15] | 0.021 | 589 |

**Table XIII.** BeviMed analysis result comparison. Shown are Bayes factors (BF) and posterior probabilities (PP) for three different priors (π).

| Gene | Phenotype | log(BF) | PP for π = 0.01 | PP for π = 0.001 | PP for π = 0.0003 | Variant impact | Mode of inheritance |
|---|---|---|---|---|---|---|---|
| ***BMPR2*** | I/HPAH | 265.762 | 1.000 | 1.000 | 1.000 | High | dominant |
| ***BMPR2*** | PAH | 265.639 | 1.000 | 1.000 | 1.000 | High | dominant |
| ***BMPR2*** | I/HPAH/PVOD/PCH | 263.481 | 1.000 | 1.000 | 1.000 | High | dominant |
| ***BMPR2*** | PH | 262.625 | 1.000 | 1.000 | 1.000 | High | dominant |
| ***BMPR2*** | young age | 149.576 | 1.000 | 1.000 | 1.000 | Moderate and high | dominant |
| ***BMPR2*** | HPAH | 149.091 | 1.000 | 1.000 | 1.000 | Moderate and high | dominant |
| ***BMPR2*** | FPAH | 147.822 | 1.000 | 1.000 | 1.000 | Moderate and high | dominant |
| ***BMPR2*** | IPAH | 144.582 | 1.000 | 1.000 | 1.000 | High | dominant |
| ***BMPR2*** | KCO higher tertile | 99.923 | 1.000 | 1.000 | 1.000 | High | dominant |
| ***BMPR2*** | middle age | 63.119 | 1.000 | 1.000 | 1.000 | Moderate and high | dominant |
| ***BMPR2*** | KCO middle tertile | 52.706 | 1.000 | 1.000 | 1.000 | Moderate and high | dominant |
| ***EIF2AK4*** | low KCO | 29.741 | 1.000 | 1.000 | 1.000 | Moderate and high | recessive |
| ***EIF2AK4*** | KCO lower tertile | 26.247 | 1.000 | 1.000 | 1.000 | Moderate and high | recessive |
| ***TBX4*** | I/HPAH | 23.783 | 1.000 | 1.000 | 1.000 | High | dominant |
| ***TBX4*** | I/HPAH/PVOD/PCH | 23.549 | 1.000 | 1.000 | 1.000 | High | dominant |
| ***TBX4*** | PAH | 23.141 | 1.000 | 1.000 | 1.000 | High | dominant |
| ***TBX4*** | PH | 22.877 | 1.000 | 1.000 | 1.000 | High | dominant |
| ***EIF2AK4*** | young age | 20.547 | 1.000 | 1.000 | 1.000 | Moderate and high | recessive |
| ***TBX4*** | IPAH | 19.990 | 1.000 | 1.000 | 1.000 | High | dominant |
| ***EIF2AK4*** | I/HPAH/PVOD/PCH | 15.718 | 1.000 | 1.000 | 1.000 | Moderate and high | recessive |
| ***ACVRL1*** | HPAH | 15.501 | 1.000 | 1.000 | 0.999 | Moderate and high | dominant |
| ***EIF2AK4*** | PAH | 15.407 | 1.000 | 1.000 | 0.999 | Moderate and high | recessive |
| ***EIF2AK4*** | PH | 15.071 | 1.000 | 1.000 | 0.999 | Moderate and high | recessive |
| ***EIF2AK4*** | PVOD/PCH | 14.441 | 1.000 | 0.999 | 0.998 | Moderate and high | recessive |
| ***AQP1*** | HPAH | 12.075 | 0.999 | 0.994 | 0.983 | Moderate | dominant |
| ***EIF2AK4*** | FPAH | 11.858 | 0.999 | 0.993 | 0.979 | High | recessive |
| ***TBX4*** | young age | 11.500 | 0.999 | 0.990 | 0.970 | High | dominant |
| ***AQP1*** | I/HPAH | 11.466 | 0.999 | 0.990 | 0.969 | Moderate and high | dominant |
| ***KDR*** | KCO lower tertile | 11.362 | 0.999 | 0.989 | 0.966 | High | dominant |
| ***AQP1*** | I/HPAH/PVOD/PCH | 11.291 | 0.999 | 0.988 | 0.964 | Moderate and high | dominant |

| | | | | | | | |
|---|---|---|---|---|---|---|---|
| ***AQP1*** | PAH | 11.047 | 0.998 | 0.984 | 0.954 | Moderate and high | dominant |
| ***AQP1*** | PH | 10.791 | 0.998 | 0.980 | 0.941 | Moderate and high | dominant |
| ***AQP1*** | FPAH | 10.023 | 0.996 | 0.958 | 0.881 | Moderate | dominant |
| ***KDR*** | old age | 9.249 | 0.991 | 0.912 | 0.774 | High | dominant |
| ***GDF2*** | I/HPAH | 9.091 | 0.989 | 0.899 | 0.746 | Moderate and high | dominant |
| ***BMPR2*** | old age | 8.913 | 0.987 | 0.881 | 0.710 | High | dominant |
| ***GDF2*** | I/HPAH/PVOD/PCH | 8.775 | 0.985 | 0.866 | 0.681 | Moderate and high | dominant |
| ***SOX17*** | young age | 8.554 | 0.981 | 0.839 | 0.631 | Moderate and high | dominant |
| ***GDF2*** | PAH | 8.478 | 0.980 | 0.828 | 0.614 | Moderate and high | dominant |
| ***ATP13A3*** | KCO higher tertile | 8.035 | 0.969 | 0.755 | 0.505 | High | dominant |

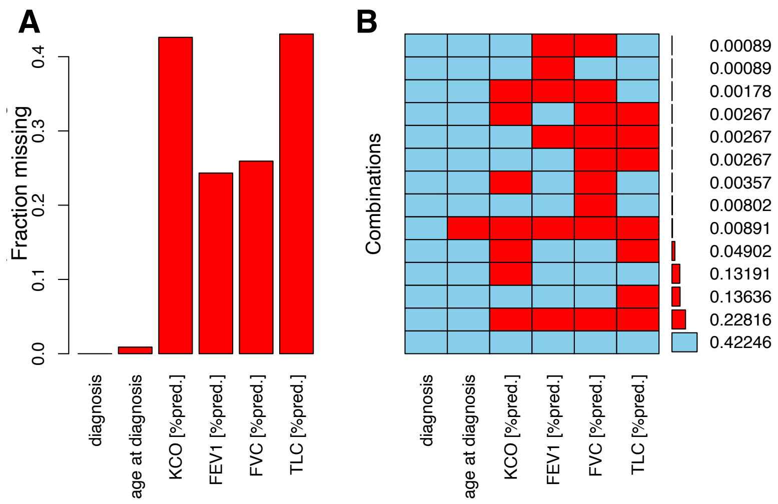


**Figure I. Summary of missing data. A**, The fraction of missing data for KCO, in comparison to diagnosis, age at diagnosis and lung function tests. **B**, The missingness pattern in KCO, in relation to diagnosis, age at diagnosis and lung function tests. KCO - transfer coefficient of carbon monoxide. $FEV_1$ - forced expiratory volume in 1 second, FVC - forced vital capacity, TLC - total lung capacity.

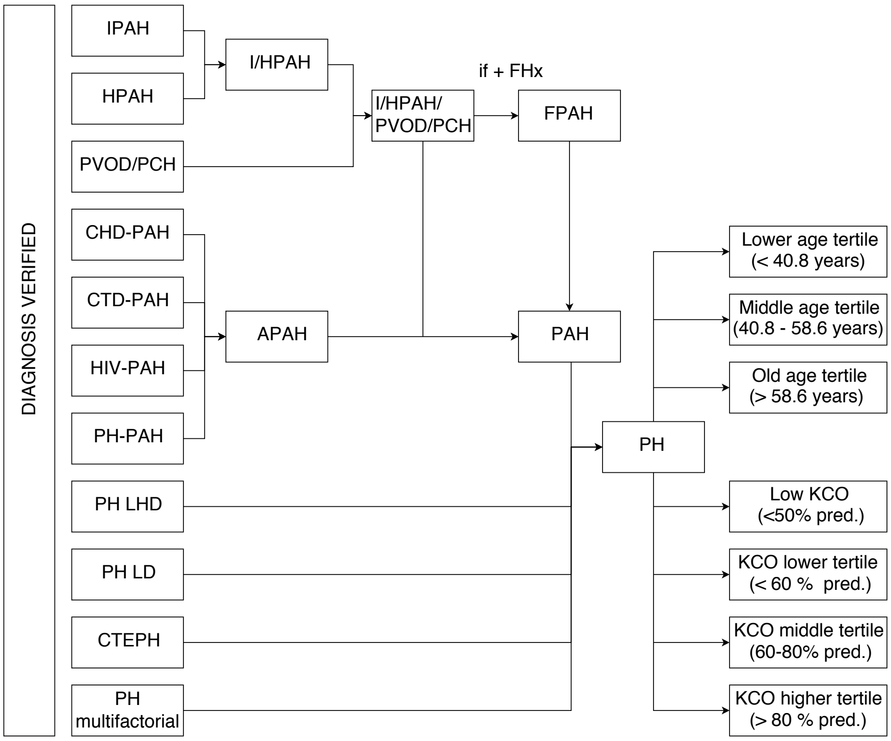


**Figure II. Flowchart describing the definition of diagnostic and phenotypic tags.** A detailed description is provided in the supplemental material. The definition of tags is listed in Table 1.

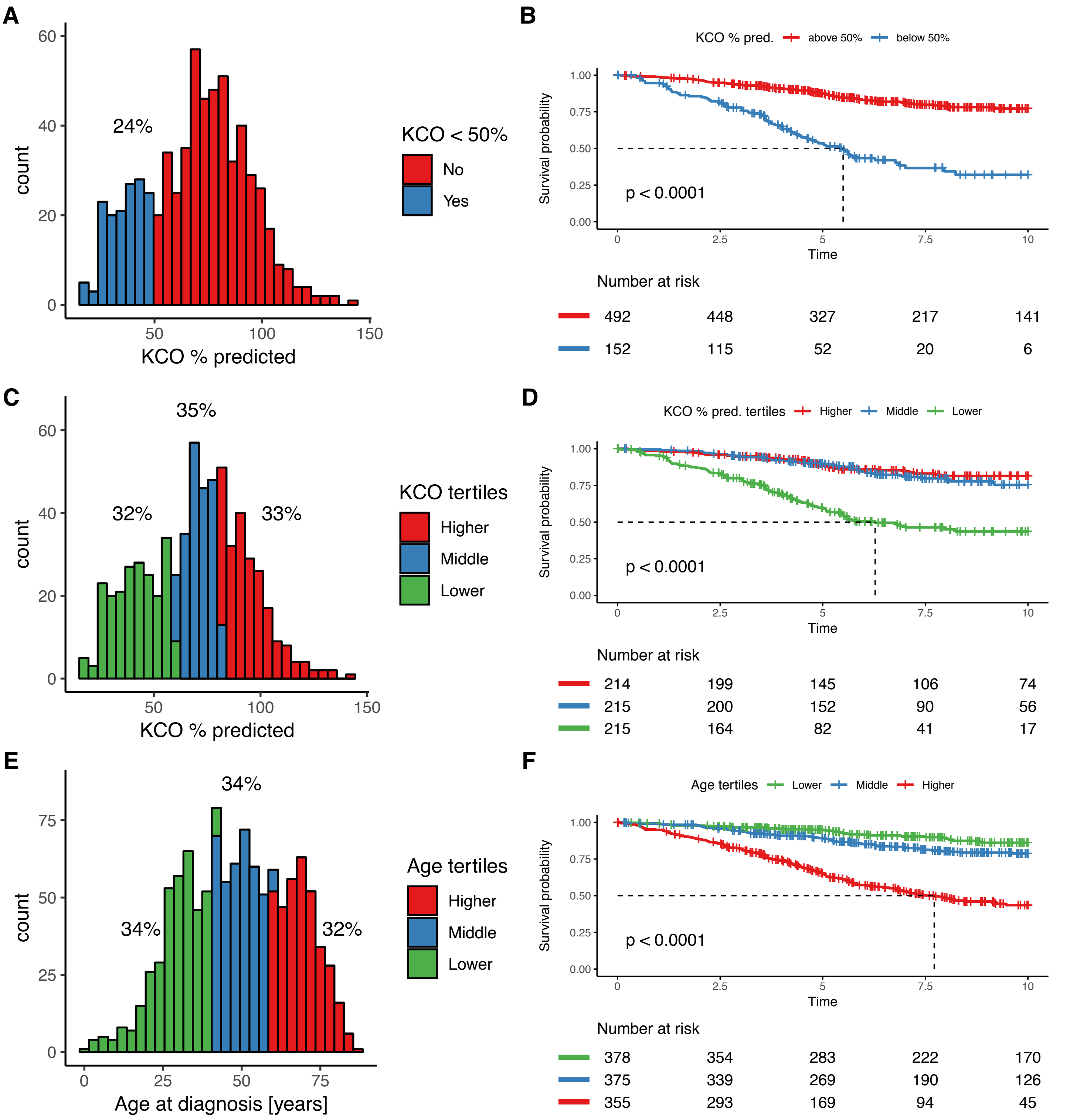


**Figure III. Characterization and survival analysis of the cohort based on transfer coefficient for carbon monoxide (KCO) and age at diagnosis.** Distribution of and Kaplan-Meier survival curves for KCO below and above the 50% predicted threshold (**A**, **B**), KCO tertiles (**C D**), and age tertiles (**E**, **F**).

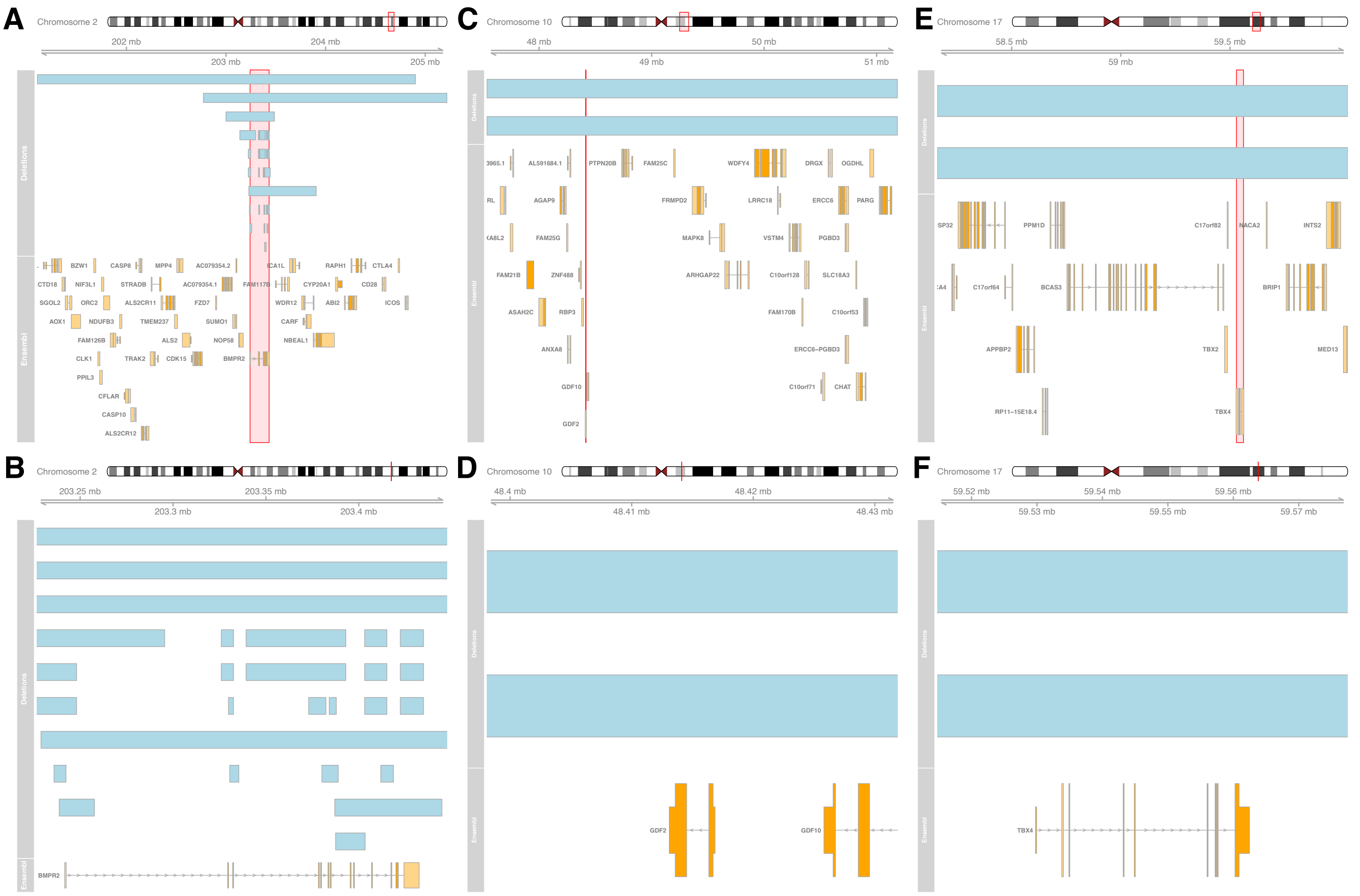


**Figure IV. Summary of large deletions identified in previously established disease genes.** Deletions are indicated by light blue boxes. The protein-coding genes, annotated in the displayed region by Ensembl (GRCh37, version 75), are depicted in the bottom panels. The affected genomic regions, with the disease gene locus highlighted in red and the magnified view focusing on the gene loci, are shown for *BMPR2* (**A**, **B**), *GDF2* (**C**, **D**) and *TBX4* (**E**, **F**).

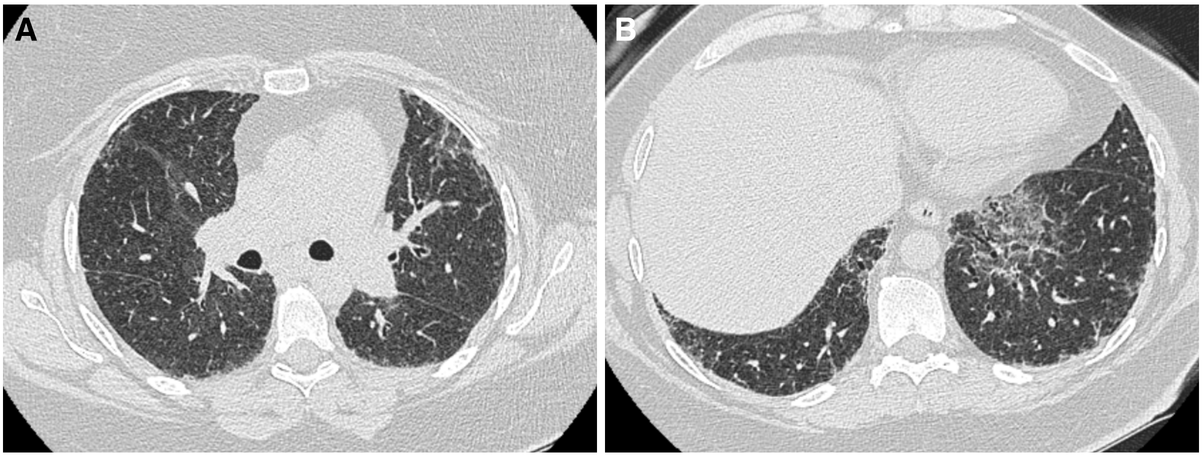


**Figure V. Chest computerized tomography (CT) scans of daughter of W000229 carrying protein-truncating *KDR* mutation. A**, The scan shows bibasal reticular ground glass changes with mild traction bronchiectasis. **B**, Upper left lobe shows sub pleural reticular ground glass changes in keeping with interstitial fibrosis. Further ground glass changes are visible in the right lung.
